# Class-wide sequence, structural, and docking-profile diversity of insect odorant receptors and structural specialization of Orco

**DOI:** 10.64898/2026.02.12.705626

**Authors:** Tianmin Zhang, Xuanxiao Yang, Yufang Fu, Wei Xue, Yifeng Zhang, Suyang Duan, Yangming Yin, Yi Guo, Chenxi Gao, Yang Liu, Gang Li, Chang Xu, Huimeng Lu

**Affiliations:** School of Life Science and Technology, Northwestern Polytechnical University, Xi’an, China; College of Life Sciences, Shaanxi Normal University, Xi’an, China

## Abstract

Insect olfaction is facilitated by a heterotetrameric odorant receptor–odorant receptor co-receptor (OR-Orco) complex, which is distinct from that of vertebrate ORs. However, extensive sequence divergence and lineage-specific expansion have complicated large-scale comparisons of OR repertoires across insects. Here, we integrated sequence similarity, structural similarity, and molecular docking to analyze ORs from 115 insect species. Across this class-wide sampling, OR repertoire size was associated with adult diet, larval diet, and habitat, but not with circadian rhythm. We developed a “trunk–branch” network framework to classify ORs according to sequence and structural similarity and used large-scale molecular docking to derive comparative docking profiles. Despite marked differences in sequence-and structural-community composition, almost all sampled orders contained ORs assigned to all six docking-profile communities. Holometabolous insects showed higher OR sequence-community diversity than non-holometabolous insects, and docking-profile composition differed between extant lineages whose order-level origins predate or postdate the end-Permian mass extinction. Most notably, we identified a highly conserved extracellular β-sheet in Orco that restricts ligand access to the binding pocket and may contribute to its reduced capacity to bind natural odorants. The emergence of Orco represents a key evolutionary transition from homomeric receptors to heteromeric OR–Orco complexes, and our results suggest that this transition may have been accompanied by pronounced specialization of the extracellular domain and binding pocket. Together, these analyses provide a class-wide framework for comparing insect OR diversity and generate testable hypotheses for future experimental and evolutionary studies.

## Introduction

Olfactory systems play a critical role in animal survival strategies, governing critical behaviors from nutrient acquisition to mate selection(Bargmann 2006; Nei et al. 2008; Kaupp 2010). Within the kingdom Animalia, insects—the most speciose metazoan group—have evolved exceptionally sophisticated chemosensory systems to interpret complex volatile landscapes(Nei et al. 2008; Kaupp 2010; Mora et al. 2011; Robertson 2019). The extraordinary sensitivity of insect olfaction to hydrophobic volatile organic compounds (VOCs) is facilitated by a heterotetrameric channel complex consisting of odorant receptors (ORs) and odorant receptor co-receptors (Orco)(Clyne et al. 1999; Vosshall et al. 1999; Sato et al. 2008). Insect ORs represent a distinct protein superfamily that originated independently of vertebrate ORs, with no shared homology between the two(Sato et al. 2008; Hansson and Stensmyr 2011). Studies have shown that this ancient structural fold has a deeper phylogenetic origin(Benton and Himmel 2023; Himmel et al. 2023). While vertebrate ORs have been extensively studied(Niimura and Nei 2005; Hughes et al. 2018; Pacalon et al. 2023; Policarpo et al. 2024), the extremely low sequence similarity among insect ORs has made resolving their internal sequence, structural, and functional differences challenging(Robertson and Wanner 2006). Extensive studies have reported advances in the olfactory recognition mechanism and OR function(Guo et al. 2021; Chang et al. 2023; Depetris-Chauvin et al. 2023; Jiang et al. 2024), which have greatly facilitated a deeper understanding of insect olfactory perception. However, a class-wide integrative framework linking OR sequence divergence, structural variation, and docking-derived binding-profile diversity is still lacking. Such a framework is needed to distinguish broadly conserved patterns from lineage-specific features and to place species-level OR diversification in a broader macroevolutionary context.

Within the class Insecta, Orcos exhibit exceptional conservation (>400 Mya evolutionary stability), whereas lineage-specific ORs display unprecedented diversity, far exceeding that of vertebrate ORs(Larsson et al. 2004; Nei et al. 2008; Brand et al. 2018; Robertson 2019; Wicher and Miazzi 2021). Although several studies have linked OR repertoire evolution to ecological adaptation in specific insect lineages, a unified class-wide comparison remains lacking(Jongepier et al. 2022; Yin et al. 2022; Singh et al. 2025). Traditional phylogeny-driven subfamily categorization, constrained by extreme cross-order sequence divergence, fails to capture the continuum of OR evolutionary innovation(Zhou et al. 2012; Zhou et al. 2015; Mitchell et al. 2020; Legan et al. 2021; Tian et al. 2022; Yin et al. 2022). Unlike vertebrate G protein-coupled ORs, insect ORs function as ligand-gated ion channels by assembling with the conserved co-receptor Orco, with recent cryo-EM structures supporting a 1:3 OR–Orco heterotetrameric model. (Jones et al. 2005; Nei et al. 2008; Kaupp 2010; Jones et al. 2011; Wang et al. 2024; Zhao et al. 2024). However, the exact in vivo stoichiometry remains to be fully resolved, and alternative arrangements such as 2:2 may also be possible(Wang et al. 2024). The mechanistic drivers of this functional divergence raise key evolutionary questions: This functional specialization raises two key questions: which structural features distinguish Orco from ligand-selective ORs, and how might these features contribute to its reduced capacity to bind natural odorants?

Identifying the binding profiles of ORs for VOCs is fundamental to understanding how insect olfactory systems perceive and adapt to their environments(McBride et al. 2014; Auer et al. 2020). To date, numerous studies have characterized the ligand-binding profiles of various insect ORs using heterologous expression systems(Hallem and Carlson 2006; Carey et al. 2010; de Fouchier et al. 2017; Guo et al. 2021; Chang et al. 2023; Jiang et al. 2024). However, when attempting to investigate the functional relevance of tens of thousands of ORs across the class Insecta at a macroevolutionary scale, experimental methods are no longer feasible due to their long timeframes and low throughput. For comparative analyses at this scale, computational approaches provide a practical means of generating hypotheses that can subsequently be prioritized for experimental testing. Although machine-learning methods remain constrained by the available training data(Boyle et al. 2013; Chepurwar et al. 2019; Caballero-Vidal et al. 2020), molecular docking offers a scalable complementary approach for comparing potential receptor–ligand interactions across large OR repertoires (Lyu et al. 2019; Comte et al. 2025). This strategy has previously been applied to mammalian OR repertoires: by employing large-scale molecular docking and OR binding profile prediction, researchers have revealed differences in olfactory perception among bats with different diets, as well as dynamic trade-offs between vision and olfaction in primates(Zhang et al. 2024; Chi et al. 2025). These results suggest that molecular docking can facilitate the exploration of macroevolutionary patterns.

In this study, we characterized large-scale patterns of OR diversification across 115 species spanning 16 insect orders. Using a framework that integrates network analysis, protein structure prediction, and molecular docking, we constructed sequence-similarity (SSN), structural-similarity (StSN), and docking-profile similarity networks (dPSN) to classify ORs from three distinct perspectives. We identified differences in OR sequence-community diversity between holometabolous and non-holometabolous insects, as well as differences in docking-profile community composition among extant lineages whose order-level origins fall on opposite sides of the end-Permian mass extinction (EPME). Across this class-wide sampling, OR repertoire size was associated with adult diet, larval diet, and habitat, but not with circadian rhythm. Moreover, the conserved β-sheet in the extracellular loop 2 (EL2) region of Orco, together with its specialized binding-pocket architecture, is associated with its reduced ligand-binding capacity and represents a prominent structural specialization of the co-receptor lineage.

## Results

### Characterization of the insect odorant receptor gene repertoire

We compiled all published insect genomes (as of December 2022), selecting the highest-quality genome assembly (contig N50 > 1 Mb) from each family as the representative species. Our study includes 115 species, featuring 8 representative species from different insect orders with previously annotated ORs(Engsontia et al. 2008; Wang et al. 2015; Zhou et al. 2015; Brand et al. 2018; Robertson et al. 2019), spanning 16 orders and 111 families. This approach ensured a comprehensive analysis across the class Insecta (Fig. 1A). To increase annotation reliability, we implemented the most stringent protein superfamily member screening strategy to date (fig. S1A). A total of 8,905 structurally intact ORs were retained for downstream analyses (Supplementary Data S2). Benchmarking against curated OR repertoires showed that the automated pipeline recovered most previously reported ORs and that stringent structural filtering produced a high-confidence set of structurally intact receptors (Supplementary Text S1, Tables S1–S3, and fig. S1B–C). For most species with comparable genome-based references, retained OR counts exceeded 70% of published values. Our analysis revealed substantial variation in OR counts across insect species. Hymenoptera exhibited the highest number of ORs and accounted for 8 of the top 10 species with the largest OR repertoires(Bonasio et al. 2010). In our OR dataset, Odonata had the fewest intact ORs, with a mean of 4 (N = 3). Hymenoptera also showed the widest range in OR counts (min: 38, max: 473, N = 23), followed by Coleoptera (min: 10, max: 271, N = 15) (Fig. 1B). Previous studies have shown that in certain insect orders and specific lineages, OR repertoire characteristics are associated with various ecological traits (McBride 2007; Liu et al. 2021; Schrader et al. 2021; Jongepier et al. 2022; Singh et al. 2025). To determine whether these relationships extend to the class Insecta, we analyzed four ecological factors: adult diet, larval diet, habitat, and circadian rhythm. Adult and larval diets represent stage-specific ecological traits, and our analysis of OR gene numbers at the genomic level aimed to reveal their relationship with long-term evolutionary pressures shaping OR gene family size. Our results revealed a significant correlation between insect OR counts and dietary preferences, both in adults (Fig. 1C, P = 1.61 × 10⁻⁶) and larvae (Fig. 1D, P = 0.03143). Habitat was also significantly associated with OR counts (P = 0.0354), with terrestrial insects exhibiting more ORs than aquatic insects do (Fig. 1E). However, there was no significant correlation between circadian rhythm and OR count (P = 0.7188, Fig. 1F). Overall, our findings suggest that insect OR counts correlate with dietary specialization (in both larvae and adults) and habitat type but are not associated with circadian rhythm. We then repeated the phylogenetic generalized least squares (pGLS) analyses using the larger of our structurally intact OR count and the published OR count, and the main ecological associations were retained (fig. S1 D-G). This indicates that the main OR-count ecological analyses are reasonably robust to moderate underestimation caused by missed ORs or strict structural filtering.

**Fig. 1.**
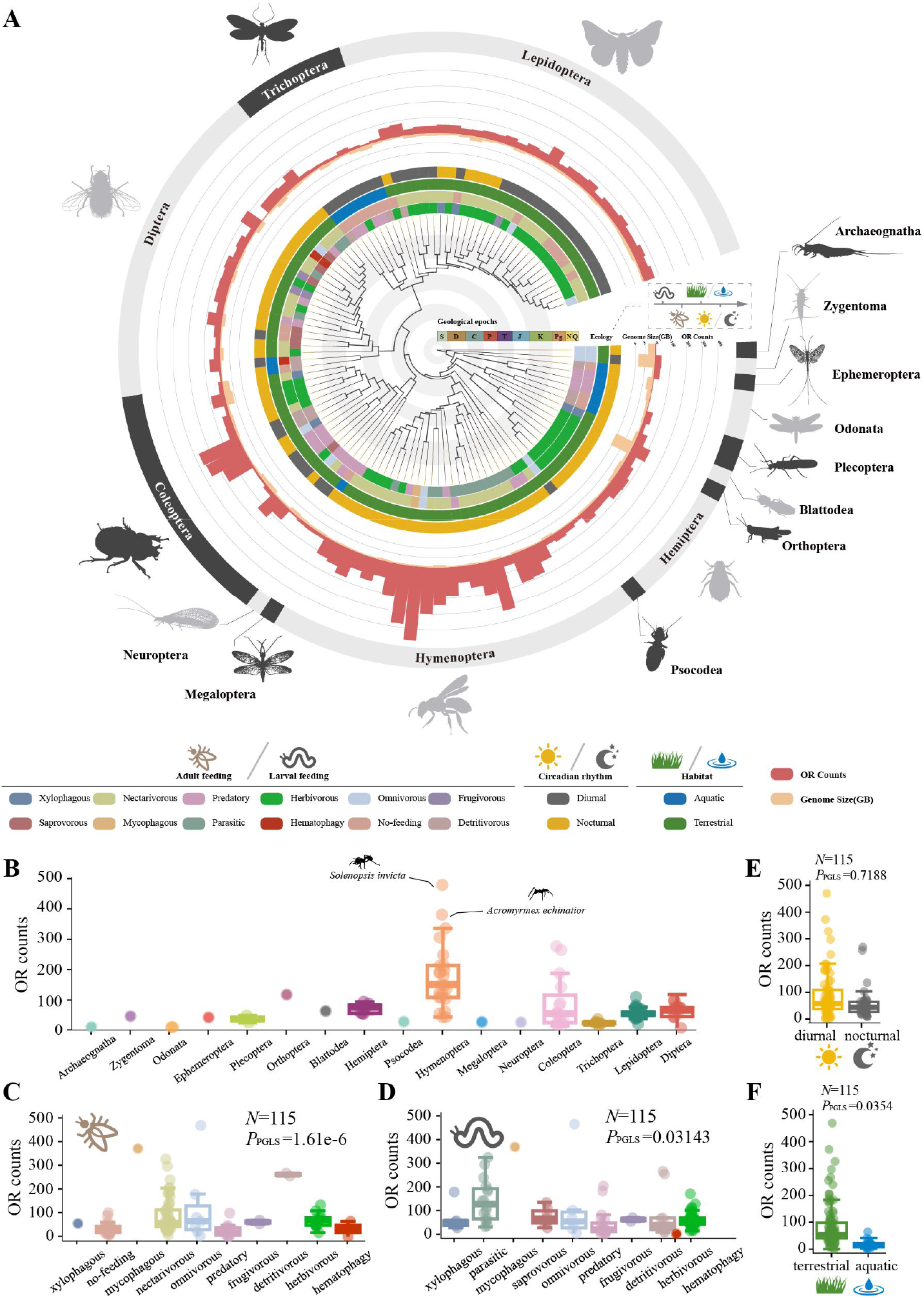
Evolutionary ecology of insect OR repertoires. (A). Phylogeny of 115 insect species (see Methods and Supplementary Data S4 for details). Genome size and the number of OR genes for each species are shown as bar plots. Ecological parameters, including larval and adult diets, habitat, and circadian rhythm, are displayed as labels with color coding corresponding to the legend below. Different geological periods are separated by shaded intervals. The tree was visualized using the iTOL(Letunic and Bork 2024). (B). Boxplot showing the number of OR genes for each insect order. (C-F). Results of two-sided pGLS tests assessing the relationship between intact OR gene numbers and various ecological parameters: adult diet (C), larval diet (D), circadian rhythm (E), and habitat (F). Dots represent OR counts of each species. N represents the counts of species. Animal silhouettes were obtained from PhyloPic.org.

### OR sequence similarity network

This study includes species representing the full phylogenetic diversity of the class Insecta, and their OR sequences exhibit substantial variation in amino acid similarity (min = 13.7%, max = 97.6%, and mean = 29.6%) (Fig. 2B). Because of the extensive sequence divergence among ORs from different insect orders, deep relationships are difficult to resolve reliably in a single class-wide phylogeny. We therefore constructed an SSN based on all-vs-all sequence comparisons as a complementary approach to characterize large-scale patterns of OR sequence similarity. In the SSN, each node represents a cluster of ORs sharing at least 60% sequence similarity, and edges are drawn between nodes if the average alignment E value < 1e-25 (fig. S2A). Nodes with a greater number of connections indicate closer evolutionary relationships.

**Fig. 2.**
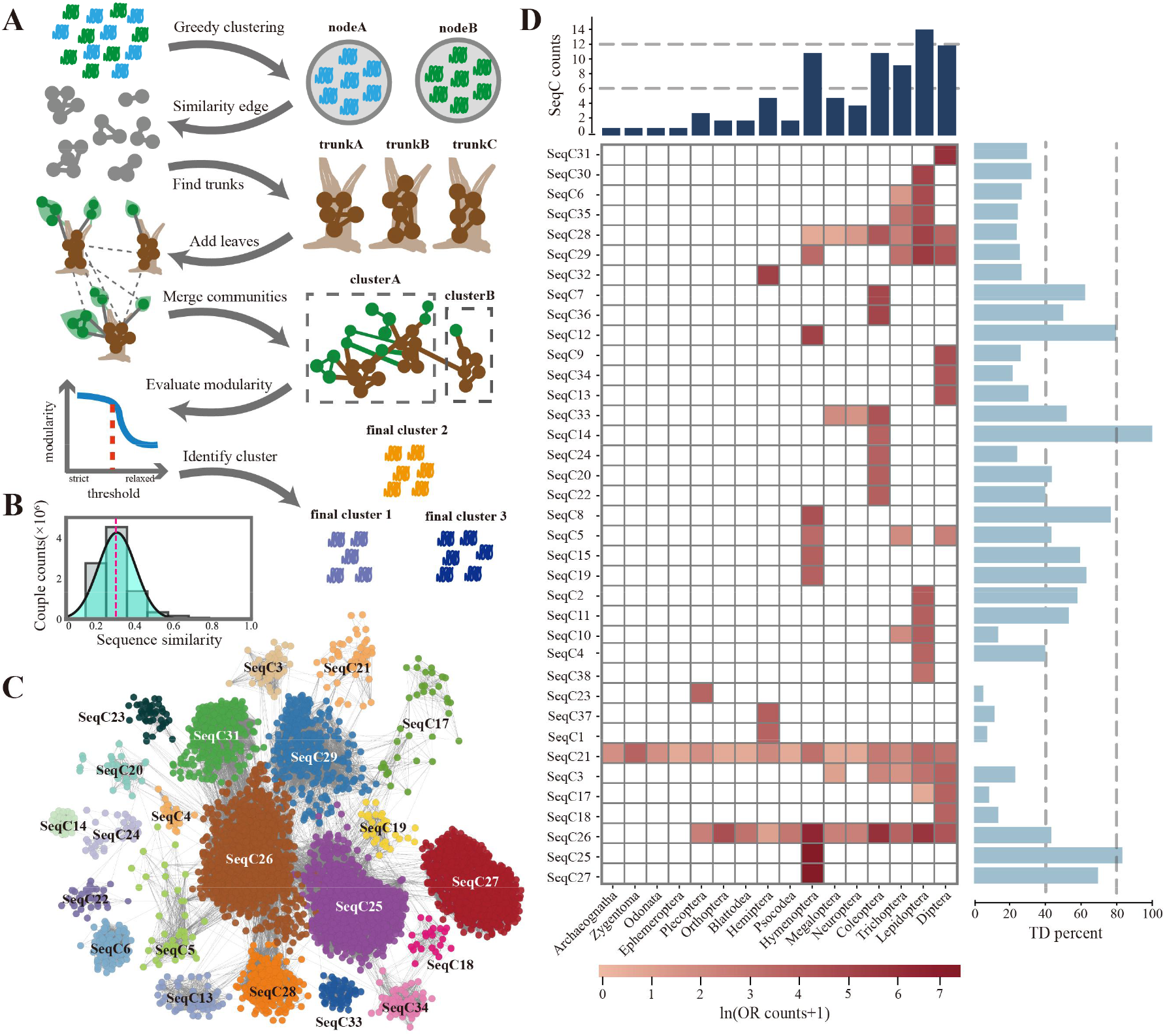
Subfamily classification of ORs based on SSN. (A). Schematic representation of the “trunk-branch” strategy pipeline for constructing the similarity network and classifying subfamilies. First, all ORs were subjected to greedy clustering based on sequence similarity to generate nodes. The average similarity between nodes was then calculated, and nodes were treated as units to identify core regions (trunks) using a strict similarity threshold. By gradually relaxing the threshold, new core regions and additional nodes (branches) emerged. After a final merging step based on defined criteria, the optimal community structure was determined using modularity as an index (see Methods). (B). Distribution of pairwise sequence similarity among all ORs. The red line indicates the mean similarity. (C). SSN of ORs constructed using the “trunk-branch” strategy. Each node represents ORs sharing >60% sequence identity. Edges represent node pairs with an average sequence alignment E-value of < 1 × 10⁻²⁵. Node colors indicate SeqC classifications. (D). Heatmap depicting the number of ORs within different SeqCs across various insect orders. Due to substantial differences in OR numbers among orders, the values are ln(count + 1)-transformed. The bar plot above the heatmap shows the number of SeqCs present in each order, while the bar plot on the right indicates the proportion of ORs within each SeqC generated by tandem duplication. Two ORs located within 5 kb of each other on the genome were defined as tandem duplicates. TD: Tandem Duplication.

Using the "trunk–branch" strategy (Fig. 2A), we identified 37 OR sequence communities (SeqCs), which together account for 94.9% of all ORs (Supplementary Data S5). Among them, 22 SeqCs demonstrated clear interaction relationships (Fig. 2C). The sequence similarity within each SeqC was predominantly between 20% and 30% (fig. S2B). By mapping previously classified *Acromyrmex echinatior* sequences using phylogenetic methods(Zhou et al. 2015), we found that most subfamilies—such as 9-exon, Orco, F, H, B, and A—were independently delineated (fig. S2C) and exhibited well-defined boundaries (fig. S2E). Our classification did not disrupt any known subfamilies and instead merged some families with minor differences (fig. S2, C and D). This comparison indicates that SSN-based communities are broadly consistent with established phylogenetic subfamilies, while capturing sequence-similarity modules rather than strictly monophyletic clades. The SSN is not intended to replace phylogenetic reconstruction, but rather to provide a complementary framework for summarizing sequence relationships among thousands of highly divergent ORs at a class-wide scale. It allows major patterns of OR sequence similarity and diversification to be visualized more clearly across the entire dataset. Phylogenetic analyses can then be used to further resolve evolutionary relationships within specific sequence communities. We further evaluated the SSN using curated lepidopteran pheromone receptors (PRs). To distinguish different PR lineages, experimentally validated classic PRs and the recently reported new PR lineage were analyzed separately(Bastin-Heline et al. 2019). Classic PRs were predominantly assigned to SeqC6 (93.9%), whereas the new PR lineage was assigned to SeqC35 (fig. S2F), supporting the hypothesis of multiple independent origins of PRs(Bastin-Heline et al. 2019). These results further support the applicability and robustness of SSN-based community partitioning.

### SeqC21 includes the Orco group as well as all ORs from *Thermobia domestica* and *Machilis hrabei*

(fig. S2G), suggesting that ORs in early wingless insects share high sequence similarity with Orco(Brand et al. 2018). In addition to SeqC21, most SeqCs are order specific, indicating substantial sequence divergence among ORs from different insect orders (Fig. 2D). SeqC26 and SeqC28 are large shared clusters that may contain cross-order orthologous relationships. In SeqC26, we identified 55 orthologous groups, four of which spanned multiple insect orders (Supplementary Data S6). OGG54 included ORs from Orthoptera, Blattodea, Trichoptera, and Hymenoptera. This group contains the locust receptors LmigOR5 and LmigOR4. LmigOR5 has been reported to bind geranyl acetone and to be associated with avoidance behavior in the migratory locust(Chang et al. 2023). The functions of the corresponding receptors in other insect orders remain to be tested, and we now present these as candidate conserved modules rather than confirmed functional orthologs. In contrast, we did not identify clear cross-order orthologous groups in SeqC28.

Lepidoptera was found to contain the greatest number of SeqCs, reflecting the highest level of OR sequence divergence. Additionally, we identified SeqC25 and SeqC27 as two exceptionally large, Hymenoptera-specific groups, comprising 39.8% and 31.9% of all Hymenopteran ORs, respectively. SeqC27 corresponds to the previously identified Hymenopteran 9-exon family (fig. S2, C and D), which originated through tandem duplication events(Engsontia et al. 2015; Zhou et al. 2015; McKenzie et al. 2016; Legan et al. 2021). Interestingly, we found that SeqC25 exhibited an even higher rate of tandem duplication than did SeqC27, suggesting that SeqC25 also expanded through extensive tandem duplication. Notably, among SeqCs with tandem duplication rates exceeding 60%, 5 of 7 are exclusive to Hymenoptera, indicating that OR lineages in this order have undergone particularly intense tandem duplication expansion compared with other insect orders(Legan et al. 2021).

### Structural diversity of insect ORs

Insect ORs fall within the "twilight zone" (10%–40% sequence identity) of sequence alignment. However, nearly all ORs remain within the "safe zone" in structural alignment (Fig. 3A).

**Fig. 3.**
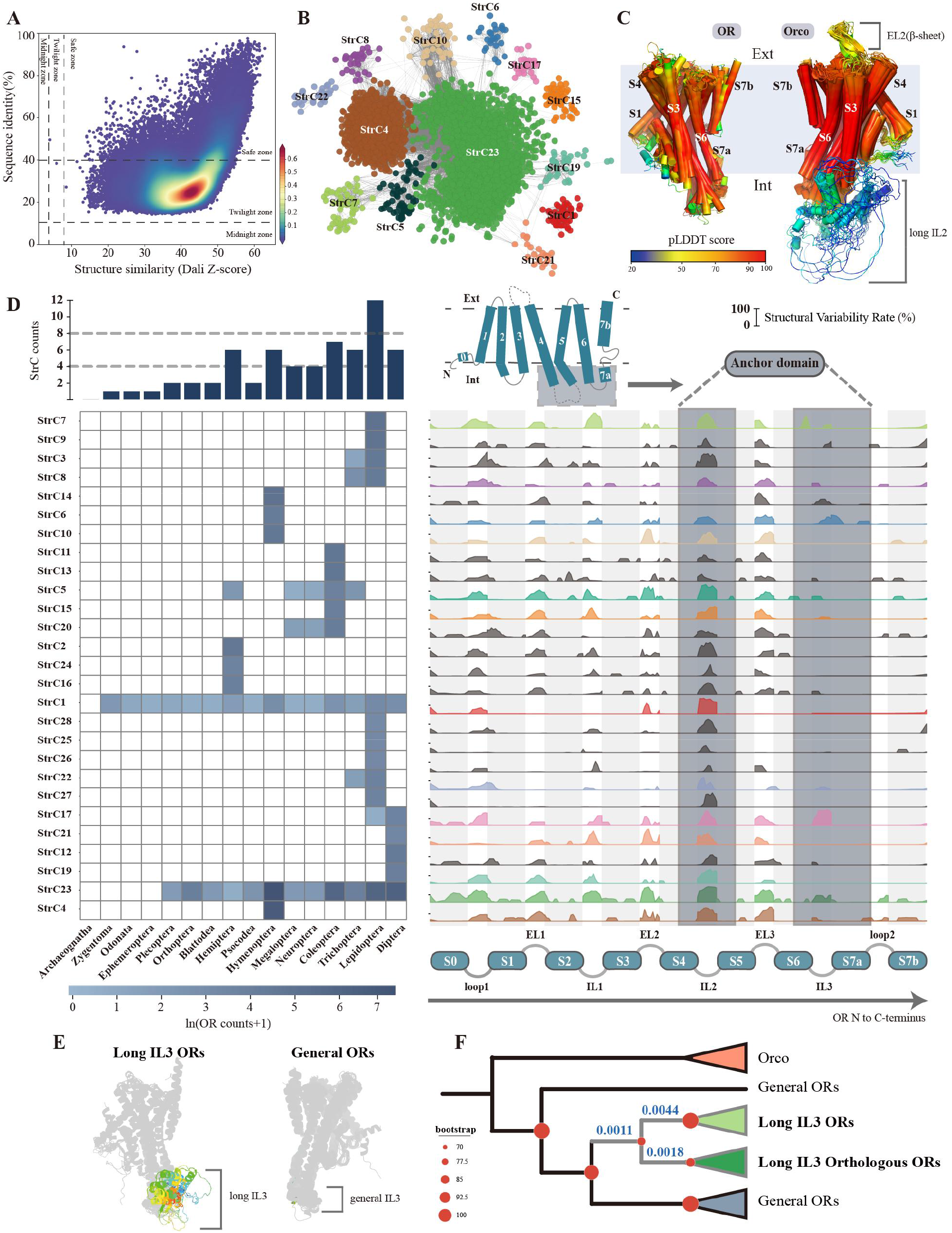
StSN and structural community diversity of ORs. (A). Sequence similarity versus structural similarity of insect ORs, assessed using pairwise comparisons with Dali. Each dot represents comparison between a pair of ORs, with color indicating point density. (B). StSN of ORs constructed using the “trunk-branch” strategy. Nodes represent ORs with pairwise Dali Z-scores >50. Edges connect nodes with an average pairwise Z-score of at least 42.9. Node colors denote StrCs classifications. (C). Structural comparison of ORs and Orco. The structures were aligned using US-align, with multiple structural alignments performed for annotated Orco and OR structures (visualized in PyMol). Colors represent pLDDT scores provided by AlphaFold2, indicating the confidence of structural modeling. (D). Visualization of differences among StrCs. The heatmap shows the number of ORs within different StrC across insect orders, with OR counts ln(count + 1)-transformed due to large inter-order variations. The bar plot above the heatmap indicates the number of StrC in each order. On the right, a ridge plot illustrates the flexibility rates of different structural regions across StrCs. Flexibility rate = 1 - conservation rate. Light gray represents transmembrane regions, white denotes loop regions, and dark gray marks the anchor domain of ORs. Colors correspond to StrCs, with gray indicating StrCs without interactions. (E). Two types of ORs in StrC17 — long IL3 ORs and general ORs — with the IL3 regions highlighted in color. (F). Phylogenetic analysis of StrC17 ORs based on amino acid sequences. Red circles indicate bootstrap values, with most ORs collapsed based on bootstrap support. Gray branches represent the clade of long IL3 ORs, and the dN/dS values are shown in blue on the corresponding evolutionary branches.

To investigate the structural diversity of insect ORs, we constructed a StSN, where each node represents ORs with Z score > 50 (from Dali(Holm 2022)). Edges were drawn between nodes if their average Z score was > 42.9 (fig. S3A). Similar to the SSN, we classified 27 structural communities (StrCs), which together accounted for 87.0% of all ORs (Supplementary Data S5). Among these, 13 StrCs exhibited distinct interaction relationships (Fig. 3B). The boundaries between StrCs were well defined (fig. S3C).

Most StrCs were also order specific (Fig. 3D). StrC1 represents the Orco group, which includes 8 *T. domestica* ORs. These ORs were previously identified as Orco-like genes through sequence analysis(Brand et al. 2018). Compared to ORs, Orco is characterized by two distinct structural features: a β-sheet in EL2 and a long loop in intracellular loop 2 (IL2) (Fig. 3C and fig. S3B). Lepidoptera presented the highest diversity of StrC types (Fig. 3D). We found that the transmembrane regions in different StrCs presented low variability, whereas the loop regions presented distinct variable patterns (Fig. 3D). In both ORs and Orcos, the IL2 region was identified as the most variable region and the only variable segment within the anchor domain. This high degree of variability may be associated with its role in stabilizing and regulating the anchoring domain(Bahk and Jones 2016; Jain et al. 2021; Wang et al. 2024; Zhao et al. 2024).

StrC23 is the only structural community present in 12 insect orders and accounts for 46.3% of all structurally intact ORs in the dataset. We found that StrC23 has a much larger binding-pocket volume than other ORs and Orco. From a structural perspective, this suggests that StrC23 receptors may have greater potential to accommodate diverse VOCs (fig. S3 D). To further investigate the evolutionary basis of this broadly shared structural feature, we examined the phylogenetic and orthology relationships among StrC23 receptors. StrC23 was divided into 156 orthologous groups, 24 of which contained ORs from more than 10 species. However, only a few of these larger groups spanned multiple insect orders: one included ORs from Hymenoptera, Diptera, and Trichoptera, while two included ORs from both Lepidoptera and Trichoptera (Supplementary Data S6). Thus, StrC23 does not represent a single conserved cross-order orthologous lineage, but instead comprises multiple phylogenetically distinct OR lineages. Its shared large-pocket architecture therefore cannot be explained solely by conservation of a single orthologous lineage and may reflect either structural convergence or retention of an ancient structural feature followed by substantial sequence divergence.

Notably, StrC17 exhibited a high degree of flexibility in the IL3 region. We observed that a subset of ORs within this group possessed unusually long IL3 regions (Fig. 3E). These ORs were distributed across 15 families from two suborders of Diptera, but they were absent from the earliest-diverging families in the Nematocera suborder, including Culicidae, Chironomidae, and Tipulidae (fig. S3, G and H). After manual inspection, annotation errors were ruled out. The construction of a maximum likelihood tree for StrC17 ORs revealed that ORs with long IL3 regions formed a distinct clade, with one subclade identified as orthologous (fig. S3F). To investigate the evolutionary pressures acting on these ORs, we calculated the dN/dS ratios for different branches (Fig. 3F, Supplementary Data S7). The results revealed that all StrC17 ORs were subject to strong purifying selection (dN/dS = 0.17). The amino acid substitution rate of ORs with long IL3 regions was two orders of magnitude lower than that of other ORs (dN/dS = 0.001–0.004), indicating intensified purifying selection.

### Structural specialization of Orco

Orco evolved from ORs in the common ancestor of Zygentoma and Pterygota (Brand et al. 2018; Thoma et al. 2019). The evolutionary drivers underlying Orco specialization remain unresolved. With the currently available early-diverging insect genomes, we could not reliably identify direct orthologous relationships among early ORs, which limits the reconstruction of ancestral OR states. The *T. domestica* ORs TdomOR1-8 are classified within Orco communities (SeqC21 and StrC1) and occupy a transitional phylogenetic position between ORs and characterized Orcos, representing the closest ORs related to Orcos(Brand et al. 2018). Because they are the extant ORs most similar to Orco in sequence, structure, and phylogenetic position, we used them as comparative proxies for candidate Orco. Comparisons between TdomOrcoFJ and the other TdomOR1–8 receptors may therefore identify structural features potentially associated with Orco recruitment and specialization in the common ancestor of Zygentoma and Pterygota. We found that TdomOrcoFJ has the smallest, least polar, and most hydrophobic binding pocket among TdomOR1-8 (Fig. 4A). Molecular docking analysis revealed that TdomOrcoFJ is less suited for VOC binding than other candidates are (fig. S4A). These findings suggest that binding-pocket specialization may have been one of the structural factors associated with the evolutionary recruitment of Orco.

**Fig. 4.**
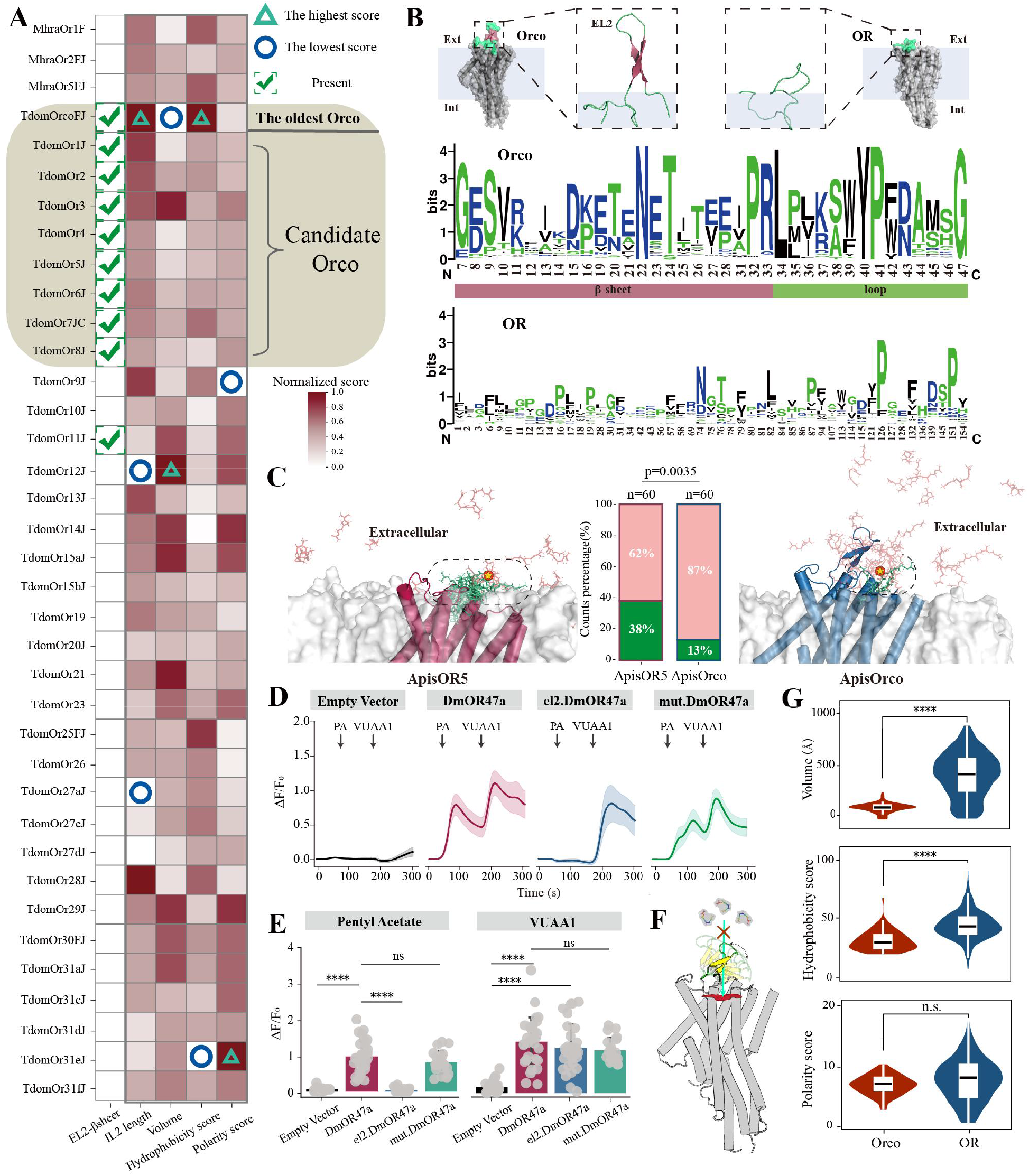
Structural features associated with Orco specialization. (A). Differences in EL2, IL2, and binding pocket properties between *M. hrabei* and *T. domestica* ORs. Only ORs with closed binding pockets were analyzed, and all scores were normalized. Circles and triangles represent the minimum and maximum scores after column normalization, respectively. The presence of an EL2 β-sheet is indicated with a check mark. (B). Differences in amino acid properties of the EL2 region between OR and Orco. Hydrophilic amino acids are shown in blue, neutral amino acids in green, and hydrophobic amino acids in black (visualized by WebLogo(Crooks et al. 2004)). The x-axis numbers represent amino acid positions from the multiple sequence alignment, with highly gapped positions removed. (C). MD simulation-based analysis of the movement trajectory of VOC (geranyl acetate) near Orco and OR. The red protein represents ApisOR5, and the blue protein represents ApisOrco. The green area indicates the probability of VOC remaining above the binding pocket, while the pink represents its probability of occupying other regions of the protein (see Methods). Statistical significance was evaluated using a Chi-Squared Test. The red pentagram indicates the initial position of geranyl acetate in the simulation. (D). Changes in relative calcium fluorescence intensity (ΔF/F0) in cells. Cells were stimulated with 100 μM of the OR47a agonist pentyl acetate (PA) at 50 s and 100 μM the Orco agonist VUAA1 at 180 s. el2.DmOR47a: OR47a mutant in which the EL2 region of OR47a was replaced with that of Orco. mut.DmOr47a: the EL2 region of wild-type DmOr47a contains an inserted short peptide. The total number of cells (n = 30) was derived from three independent replicate experiments. The shading indicates the 95% confidence interval. (E). Relative calcium fluorescence intensity in cells following stimulation with PA and VUAA1, respectively. n=30. Statistical significance was determined using Tukey’s HSD test. (F). Schematic illustration of the mechanism by which the β-sheet structure in the EL2 region of Orco influences VOC proximity to the binding pocket. (G). Comparative analysis of binding pocket parameters (volume, hydrophobicity score, and polarity score) between Orco and ORs across all studied insect species. Statistical significance was determined using a two-sided t-test. **** P < 0.0001; n.s., not significant.

The reasons behind the loss of Orco function remain unclear, although its conserved binding pocket may act as a molecular sieve(Pacalon et al. 2023). We found that *M. hrabei* ORs lack the EL2 β-sheet but have a long IL2 loop region. Similarly, most *T. domestica* ORs also have a long IL2 loop region, whereas only TdomOR1-8 retains the EL2 β-sheet (Fig. 4A). The long IL2 loop was present in ORs from early-diverging insects as well as in candidate Orco and therefore does not provide a distinguishing structural criterion for candidate Orcos. In contrast, the EL2 β-sheet was consistently present in TdomOR1–8 and highly conserved across characterized Orcos. This distribution suggests that the EL2 β-sheet may have emerged as a key structural innovation during Orco specialization and subsequently become conserved in the Orco lineage.

The formation of the EL2 β-sheet may be associated with the loss of Orco function. We observed that the EL2 region of Orco is distinctly divided into two parts: the β-sheet region and the transmembrane loop region (Fig. 4B). In the β-sheet region, the proportion of hydrophilic amino acids is 47.8%, whereas it is 14.2% in the transmembrane loop region and 18% in ORs. Overall, across all the insects, the β-sheet region of Orco EL2 presented a significantly greater proportion of hydrophilic amino acids than did the ORs (fig. S4B), facilitating its stable presence in the extracellular aqueous environment. The extracellular port is the main pathway through which ligands enter the binding pocket(Renthal and Chen 2022; Pacalon et al. 2023). Using molecular dynamics (MD) simulations to analyze VOC movement outside the membrane, we found that the presence of the β-sheet in EL2 region significantly reduced the likelihood of VOCs reaching the binding pocket (Fig. 4C, fig. S4C and Movie S1). Using Drosophila OR47a as an example, we demonstrated through calcium imaging experiments that replacing the EL2 region of OR47a with that of Orco resulted in a mutant OR47a (el2. DmOR47a) that exhibited markedly reduced calcium responses to pentyl acetate (PA) (Fig. 4, D and E and fig. S4D). Additionally, we inserted a non– β-sheet short peptide into the EL2 region of OR47a, and the results showed that its function remained unaffected (Fig. 4, D and E and fig. S4D). This finding suggests that the β-sheet acts as a spatial barrier, limiting VOC access to the binding pocket as a primary physical gate (Fig. 4F). Furthermore, we observed that Orco has a smaller and less hydrophobic binding pocket than ORs do (Fig. 4G). These characteristics are unfavorable for VOC entry into the binding pocket, representing the key chemical factors contributing to Orco’s loss of natural VOC-binding function.

### Classification of ORs based on docking-profile similarity

Currently, molecular docking and virtual screening have been successfully applied to the high-throughput prediction of OR binding profiles in mammals, facilitating the exploration of macroevolutionary patterns in predicted olfactory recognition(Zhang et al. 2024; Chi et al. 2025). To better compare the potential binding profiles of insect OR repertoires and classify them accordingly, we collected 14,499 VOCs from 4 molecular libraries (including pheromones(Li et al. 2023), fragrances (thegoodscentscompany.com), food odors(Chi et al. 2025), and theoretical odors(Mayhew et al. 2022)). These VOCs were categorized into 9 functional groups (Fig. 5A). Although terpenoids are important in insect chemical ecology, they are distributed across multiple regions of odor space (fig. S5C). To reduce the docking instability that could result from excessive VOC subdivision, we did not treat terpenoids as a category parallel to the nine basic functional-group categories. We used this comprehensive VOC library to systematically evaluate the docking-derived binding profiles of ORs, aiming to characterize the potential ligand-binding diversity encoded by insect OR repertoires, even though some of these compounds may never be encountered by insects in nature.

**Fig. 5.**
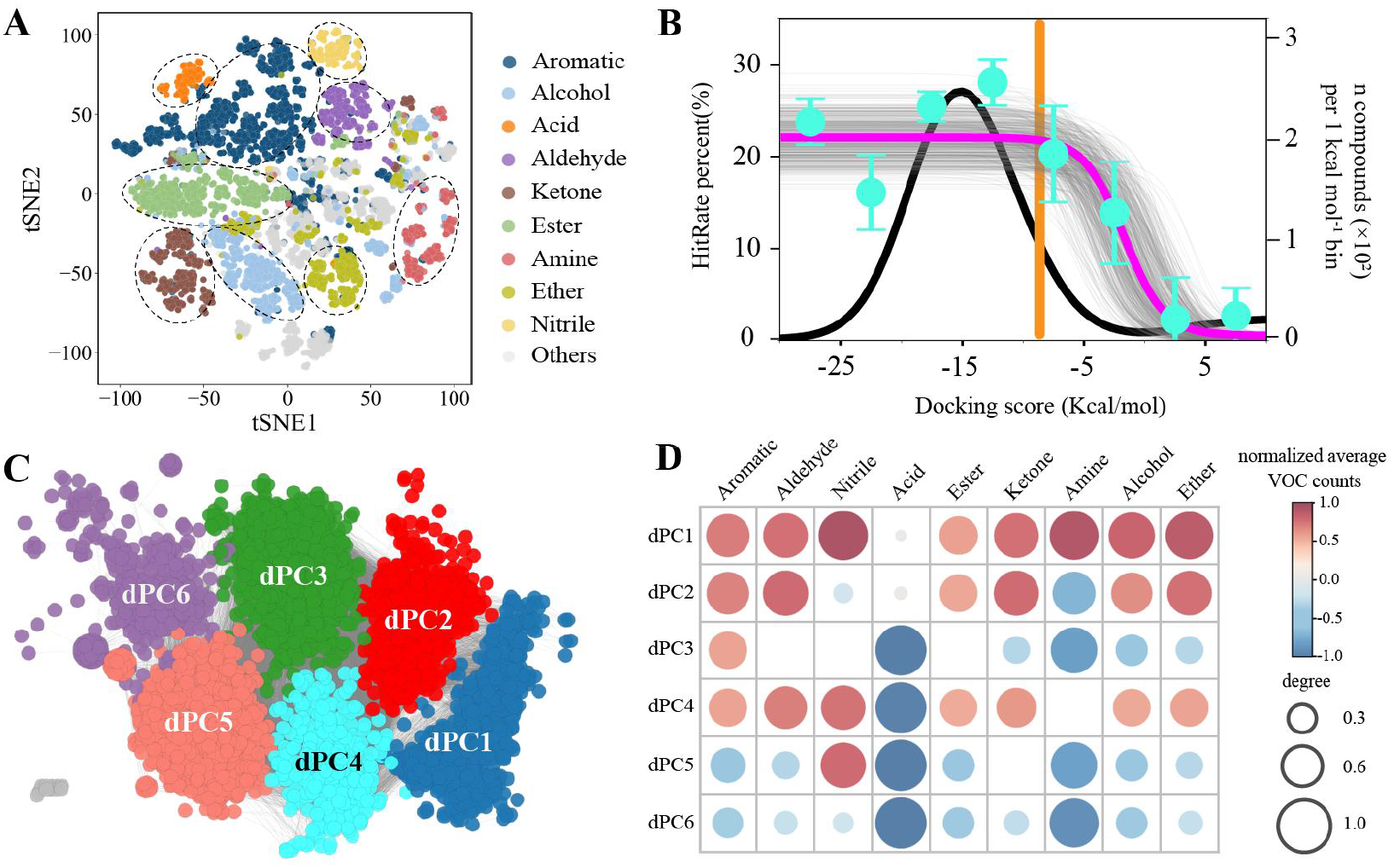
Classification of insect OR docking-profile communities. (A). 2D odor space based on molecule physicochemical properties. Each dot represents an odor molecule, with different colors indicating different functional groups. (B). Relationship between hit rate and docking score, inferred using Bayesian statistics. The black curve shows the distribution of hit rates and docking scores across all OR-VOC docking results. The top plateau has a hit rate of 22%, while the bottom plateau has a hit rate of 0%. Cyan dots represent the mean hit rate ± s.e.m. for each docking score interval. The orange line indicates the peak hit rate at a docking score of -8 kcal/mol. The gray curve represents posterior distributions from Bayesian inference, with n = 500. (C). dPSN and dPC of ORs. Nodes represent ORs with pairwise PCCs > 0.6 based on docking scores. Edges represent node pairs with an average PCC ≥ 0.48. (D). Labels for the recognition of different functional group small molecules by each dPC. Red indicates a tendency to recognize a higher number of odor molecules from a particular functional group, while blue indicates to recognize fewer. Circle size represents the degree of this tendency.

We performed molecular docking for all insect ORs using these VOCs to generate their docking-derived binding profiles (fig. S5A). A long-standing challenge in molecular docking is how to predict binding likelihood on the basis of ranking scores. To address this issue, we modeled a theoretical "hit rate" curve using docking scores and experimental datasets from Drosophila(Hallem and Carlson 2006), mosquitoes(Carey et al. 2010), and moths(de Fouchier et al. 2017) (Fig. 5B). Using a Bayesian framework with prior probability distributions, we determined an informative docking-score range for insect ORs. A threshold score of -8 kcal mol-1 was established to maximize data retention while maintaining a reliable hit rate (Fig. 5B). In addition, our single-taxon hit-rate sensitivity analysis showed similar trends for Drosophila, Anopheles, and moths (fig. S5D-F). Because the locust OR repertoire had an extremely low positive rate, we did not include the locust experimental dataset in the final threshold assessment (fig. S5 G and H, see Supplementary Text S2 for details). Although this threshold enriches experimentally responsive OR-VOC pairs, the resulting hit rate is not sufficient to support deterministic inference for individual receptor-ligand pairs. Therefore, subsequent analyses were interpreted at the OR repertoire level as docking-derived comparative patterns rather than as experimentally validated OR response spectra.

We constructed a dPSN for ORs (Fig. 5C). In this network, each node represents ORs with a docking-profile Pearson correlation coefficient (PCC) greater than 0.6, and nodes with an average PCC greater than 0.48 are connected by edges. We identified 6 docking-profile communities (dPCs), representing 72.5% of the ORs (fig. S5B, Supplementary Data S5). It should be noted that dPCs represent docking-profile communities rather than experimentally confirmed functional classes. We subsequently quantified the number of VOCs from different functional groups meeting the docking threshold for each dPC (fig. S5I) and assigned docking-derived binding labels on the basis of their relative enrichment compared with the background (Fig. 5D). These labels summarize the predicted breadth of VOC functional-group binding for each dPC. To evaluate the extent to which these docking-derived labels were consistent with available experimental data, we calculated the area under the receiver operating characteristic curve (AUC) by comparing experimental OR response data with the corresponding docking-derived labels. For most functional group predictions, an AUC exceeding 0.75 was achieved, suggesting that these labels capture broad directional tendencies in available experimental data (fig. S5J). dPC1 showed a relatively broad docking profile across VOC functional groups, whereas dPC6 showed a narrower docking profile (Fig. 5D). Additionally, dPC3 showed relatively stronger predicted binding toward aromatics, whereas dPC5 showed relatively stronger predicted binding toward nitriles.

Unlike SeqC and StrC, almost all insect orders contain all dPCs without exhibiting specificity (fig. S5K). dPC1, characterized by a relatively broad docking profile, accounted for the largest proportion of ORs in most orders, particularly Psocodea, Diptera, and Lepidoptera (fig. S5L). In contrast, dPC6, characterized by a relatively narrow docking profile, represented only a small proportion of ORs across the sampled species.

### Comparative patterns of insect OR diversification across evolutionary periods

Although the sequence evolution of the insect OR gene family and functional diversity among different orders and species have been reported multiple times(Brand and Ramirez 2017; McKenzie and Kronauer 2018; Legan et al. 2021; Chang et al. 2023), these studies reveal only a partial view rather than the broader patterns of insect OR sequence, structural, and docking-profile diversity. To characterize large-scale diversification patterns of insect ORs, we integrated SeqCs, StrCs, and dPCs. The results revealed strong congruence between SeqCs and StrCs, with both communities including nearly all dPCs (Fig. 6 and fig. S6). This pattern suggests that sequence and structural diversification are more closely coupled, whereas docking-profile variation shows a weaker correspondence with either level.

**Fig. 6.**
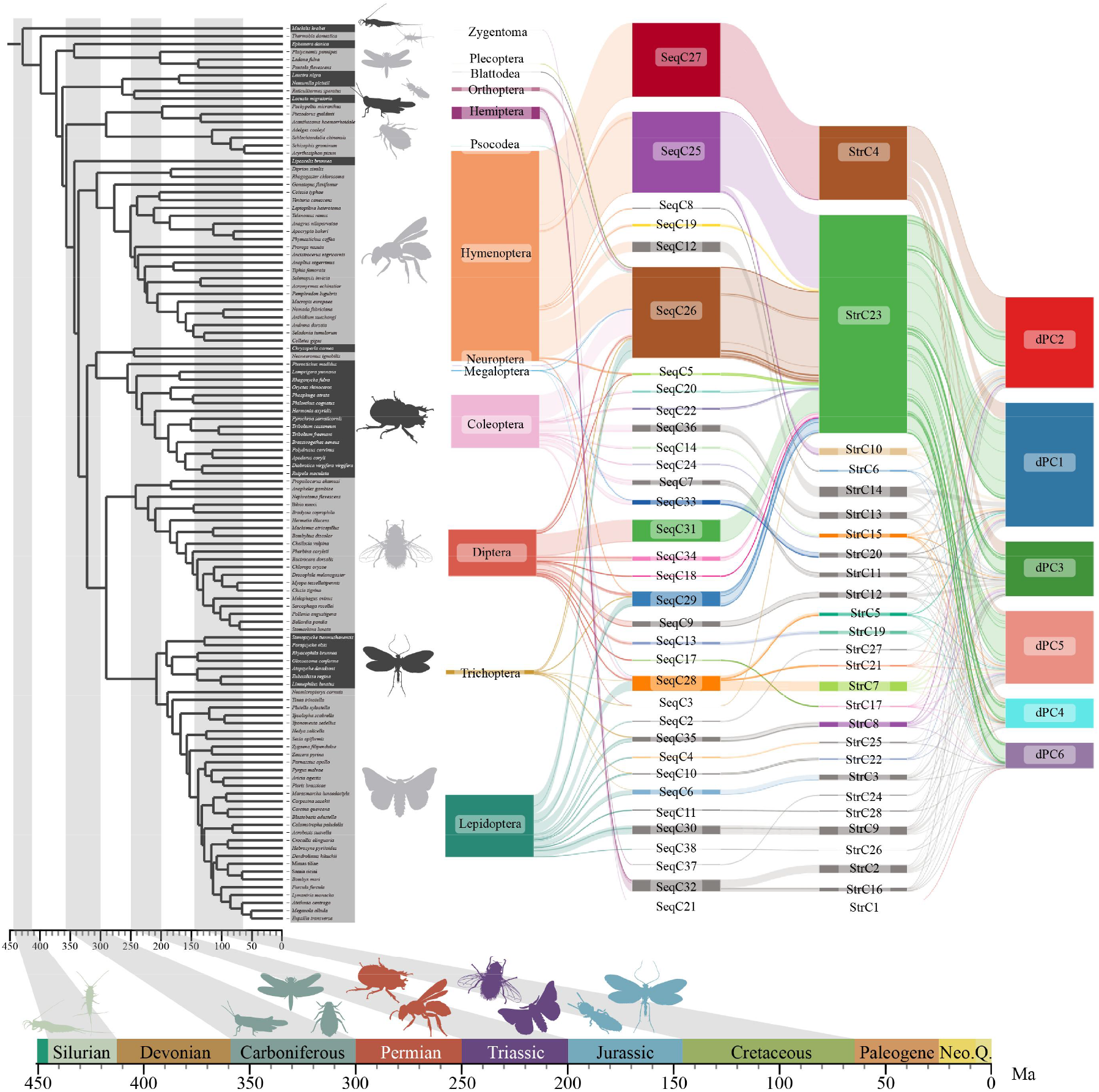
Relationships between OR sequence, structure and docking-profile communities across insect orders. The phylogenetic tree follows the structure from Fig.1. On the right, the mapping relationships between all insect OR SeqCs, StrCs and dPCs are shown, considering only ORs classified into communities. Time nodes for different geological periods are based on previous study(Misof et al. 2014) and are separated by gray shading. The timeline beneath the tree marks the origin of various insect orders. Animal silhouettes were obtained from PhyloPic.org.

Additionally, extant holometabolous insects showed higher OR sequence-community diversity than non-holometabolous insects (fig. S7). Because total SeqC counts may be affected by unequal species sampling and OR numbers, we added normalized analyses. First, we calculated SeqC numbers at the species level and found that holometabolous insects contained more SeqCs on average (fig. S8A). Second, because OR number was significantly correlated with SeqC count (P=0.001035), we performed equal-OR random sampling between holometabolous and non-holometabolous insects. Under the same OR sampling depth, holometabolous insects still covered more SeqCs, indicating higher OR sequence-community diversity after controlling for OR number (fig. S8B). Therefore, this result supports a genuine difference in SeqC diversity rather than a difference driven by unequal species sampling or OR number. The higher SeqC diversity observed in Holometabola was retained when the comparison was restricted to Neoptera, indicating that this pattern is not simply a restatement of previously reported Paleoptera-Neoptera differences (fig. S9). The number of SeqCs increased with deeper divergence among insect orders, suggesting gradual accumulation of OR sequence variation during insect evolution (fig. S8C). Despite the limited SeqC diversity in non-holometabolous insects, all six dPC types were represented in these extant lineages, suggesting that the major docking-profile community types may have arisen relatively early in insect evolution.

The EPME was a significant geological event that resulted in the extinction of most species and caused drastic environmental changes(Close et al. 2020; Jouault et al. 2022). In the extant species sampled here, we detected differences in dPC composition between lineages whose order-level origins fall before and after the EPME (fig. S8D). Lineages originating after the EPME showed a greater proportion of dPC1, which was characterized by a broader docking profile, and a lower proportion of dPC2 (fig. S8E). This pattern is consistent with previously reported data that the proportion of broad-tuned ORs in Locusta is much smaller than that in Drosophila and Anopheles (Chang et al. 2023). These observations should be interpreted as a docking-derived comparative pattern among extant lineages rather than direct evidence of ancestral functional changes across the EPME.

## Discussion

In this study, we integrated sequence, structural, and docking-profile analyses to characterize large-scale patterns of insect OR diversification. Across the sampled insect species, OR repertoire size was associated with several ecological traits. Sequence and structural communities showed a close correspondence, whereas docking-profile communities were less closely associated with either classification. Differences in dPC composition among extant lineages whose order-level origins predate or postdate the EPME raise the hypothesis that these differences may be associated with ecological changes surrounding this event. Within the “conserved chassis–diversified sensors” model, we further examined Orco specialization and identified the conserved EL2 β-sheet and distinctive binding-pocket properties as structural features that may contribute to reduced odorant-binding capacity.

The long-IL3 OR orthologous group includes *Drosophila melanogaster* DmOr13a and *Bactrocera dorsalis* BdorOR13a, both of which respond to 1-octen-3-ol and belong to a group of dipteran fly OR homologs associated with oviposition behavior(Kreher et al. 2008; Liu et al. 2023; Xu et al. 2023). Modeling of these homologous ORs showed that they also contain a long IL3 region. We therefore propose that long-IL3 ORs correspond to these fly homologs that recognize 1-octen-3-ol. However, 1-octen-3-ol has different behavioral functions in different species: it acts as an attractant in blood-feeding mosquitoes and tsetse flies(Hall et al. 1984; Kline et al. 2007), whereas in flies it is usually associated with oviposition regulation(Kreher et al. 2008; Liu et al. 2023). Mosquito receptors known to detect 1-octen-3-ol, such as AgOR8, AaOR8, TaOR8, and CquiOR118b(Lu et al. 2007; Xu et al. 2015; Dekel et al. 2016; Frunze et al. 2024), do not contain a long IL3 region (fig. S3 E). This suggests that the long IL3 structure in fly homologs may be related to the oviposition-associated role of 1-octen-3-ol in flies, which requires future experimental validation.

Despite substantial differences in OR sequence and structural community composition, nearly all sampled insect orders contained representatives of all major docking-profile communities. SeqCs and StrCs showed a closer correspondence with each other than either did with dPCs, suggesting that sequence and structural diversification are more tightly coupled, whereas variation in docking profiles is less directly associated with either level. Thus, distinct sequence and structural repertoires can converge on broadly similar distributions of docking-profile communities across insect orders. This pattern suggests that diversification in sequence and structure does not necessarily translate into equivalent divergence in predicted ligand-binding profiles. We also detected differences in dPC composition between extant lineages whose order-level origins predate or postdate the EPME. In particular, lineages whose order-level origins postdate the EPME had a higher proportion of dPC1, a community characterized by a relatively broad docking profile. The Permian–Triassic transition was accompanied by extensive ecological disruption and major restructuring of terrestrial ecosystems, including substantial changes in plant communities and insect–plant interactions (Grauvogel-Stamm and Ash 2005; Benton 2016; Vajda et al. 2020; Lu et al. 2021; Jouault et al. 2022). These environmental changes raise the hypothesis that ecological turnover surrounding the EPME may have contributed to differences in OR docking-profile composition among insect lineages. However, this interpretation remains hypothetical. Because rapid OR evolution and limited cross-order orthology preclude reliable reconstruction of deep ancestral repertoires, the EPME-related pattern should be viewed as a descriptive comparison among extant lineages. Further validation will require broader taxon sampling, ancestral reconstruction, and experimental testing of receptor tuning breadth.

Previous studies have proposed an evolutionary transition from ancestral chemoreceptors to the insect OR–Orco system, and our results provide structural insight into the specialization of Orco within this framework.(Brand et al. 2018; Thoma et al. 2019). The ancestral insect recruited GRs to perform olfactory functions, giving rise to the earliest ORs(Robertson et al. 2003; Frank et al. 2024; Mitchell et al. 2026). These early ORs retained the homotetrameric architecture of GRs(Del Marmol et al. 2021). However, the sensitivity requirements of olfaction and gustation differ(de Fouchier et al. 2017; Pask et al. 2017; Slone et al. 2017; Dweck and Carlson 2023; Wang et al. 2024; Zhao et al. 2024). This issue appears to have been first addressed in the ancestor of Zygentoma and Pterygota by reducing the number of functional subunits while maintaining the tetrameric structure, thereby increasing sensitivity(Brand et al. 2018; Thoma et al. 2019; Wang et al. 2024; Zhao et al. 2024). Two structural features are consistently conserved among Orcos: the EL2 β-sheet and the highly specialized binding pocket. The β-sheet physically prevents VOCs from accessing the binding pocket, whereas the chemically restrictive nature of the binding pocket prevents VOCs from penetrating deeply. Orco exhibits strong sequence conservation, ensuring that these chemical and physical barriers are effectively preserved. Sequence similarity is much lower in the insect OR family than in the GR family, which may be related to the emergence of Orco. This division between a conserved co-receptor and diversified ligand-selective subunits has been proposed to provide a stable structural framework within which OR diversification can occur(Zhao et al. 2024). However, we also identified several ligand-selective ORs containing an Orco-like EL2 β-sheet. Whether this structural similarity reflects shared ancestral features, convergent structural evolution, or other evolutionary processes remains unresolved (fig. S4, E and F).

Overall, we characterized the relationships among sequence, structure, and docking profiles in insect ORs, identified structural features potentially associated with Orco specialization, and detected class-wide associations between OR repertoire size and ecological traits. The resulting dataset and similarity-based community framework provide a resource for insect OR research, while the docking-profile patterns offer testable hypotheses for future experimental and evolutionary studies.

## Materials and Methods

### Genome data

We compiled data from all published insect genomes available in the NCBI public database as of December 2022. Genomes with a contig N50 > 1 Mb were considered reliable for OR annotation. When multiple high-quality genomes were available within a single insect family, we selected the genome with the highest assembly quality as the representative. After filtering, 107 insect genomes were included: Odonata (2), Plecoptera (2), Blattodea (1), Hemiptera (6), Psocodea (1), Hymenoptera (22), Megaloptera (1), Neuroptera (1), Coleoptera (14), Trichoptera (7), Lepidoptera (30), and Diptera (20). Detailed genome information is provided in the Supplementary Data S1. To maximize phylogenetic diversity, we added eight representative species from different orders with annotated ORs (*Machilis hrabei*(Brand et al. 2018), *Thermobia domestica*(Brand et al. 2018), *Ladona fulva*(Brand et al. 2018), *Ephemera danica*(Brand et al. 2018), *Locusta migratoria*(Wang et al. 2015), *Acyrthosiphon pisum*(Robertson et al. 2019), *Acromyrmex echinatior*(Zhou et al. 2015), and *Tribolium castaneum*(RRID:SCR_010440)(Engsontia et al. 2008)). In total, our study includes 115 insect species, covering 16 orders and 111 families.

### Species Phylogenies

To construct the phylogenetic tree used in this study, we merged existing phylogenies across different insect orders(Misof et al. 2014). For relationships among orders within Insecta, we referred to previously published phylogenetic frameworks. To ensure broad taxonomic representation, we included 16 insect orders in our analysis. Intra-order relationships were obtained from insectphylo.org and other published species trees covering various insect taxa(Chesters 2017). For species with available genomes that were not included in existing phylogenies, we inferred their phylogenetic placements based on family-level information. Divergence times among orders were obtained from TimeTree.org(Kumar et al. 2022) and calibrated using MCMCtree(Rannala and Yang 2007). The resulting phylogeny includes 115 insect species spanning 16 orders.

### Ecological Trait Data of Insect Species

Given the vast diversity and ecological complexity of insects, and the lack of a comprehensive authoritative database, we manually retrieved ecological trait data from published literature for each species. We characterized insect ecology based on four aspects: larval diet, adult diet, habitat, and circadian rhythm. For dietary traits, we classified diets into 12 types: herbivorous, xylophagous, hematophagy, mycophagous, non-feeding, omnivorous, predatory, saprovorous, frugivorous, nectarivorous, detritivorous, and parasitic. For habitat, following previous classification criteria, species were assigned as either terrestrial or aquatic(Dijkstra et al. 2014). Insects that live entirely or have at least one life stage in water were considered aquatic; those that are strictly land-dwelling throughout their life cycle were considered terrestrial. Circadian rhythm was classified based on adult activity periods as either diurnal or nocturnal. The ecological classification and corresponding literature sources for each species are provided in the Supplementary Data S3.

### OR gene mining and structural filtering

All OR and Orco protein sequences were downloaded from NCBI, supplemented with ORs reported in the paper, yielding a total of 16,176 ORs. Redundancy and data cleaning were then performed using CD-HIT(Li and Godzik 2006; Fu et al. 2012) v4.8.1 to remove highly similar sequences with >90% identity, resulting in 6,029 non-redundant ORs used as queries. We used Exonerate(Slater and Birney 2005) v2.4.0 to search whole-genome data and identify potential exon regions. To improve annotation efficiency, genome contigs were split into 20 Mb segments, with 20 kb flanks added at breakpoints to avoid truncating ORs. Next, InsectOR(Karpe et al. 2021) was used for OR gene annotation.

Given that multi-exon gene annotation from genome can be error-prone, we applied the strictest structural filtering strategy to enhance annotation reliability (Fig.S1d). All annotated ORs were modeled using AlphaFold2(Jumper et al. 2021). Structural alignment against the cryo-EM structure of MhOR5 (PDB: 7LIC) was performed using Dali(Holm 2022), and ORs with Z-scores <20 were excluded. We further manually removed sequences lacking at least seven transmembrane domains. In total, 8,905 ORs with complete and credible structural models were retained for downstream analyses. This rigorous approach eliminates structurally compromised ORs that may appear intact at the sequence level, thereby increasing the likelihood of identifying ORs with functional physiological and biochemical properties.

### Phylogenetic comparative analyses

The function pGLS of the R package caper was used to perform all phylogenetic generalized least squares models presented in this study. Phylogenetic manipulations were conducted using the R packages phytools (RRID:SCR_015502) and ape. Phylogenetic trees were visualized by iTOL(Letunic and Bork 2024) (https://itol.embl.de/). Animal silhouettes used in this study were retrieved from http://phylopic.org/.

### "Trunk-Branch" strategy

Most existing insect ORs have arisen through gene duplication events(McKenzie and Kronauer 2018; Legan et al. 2021). Initially, these duplicated OR sequences were highly similar, forming the “trunk” of core communities. Over time, mutations led ORs to diverge from their original sequences and functional features, forming the “branches” of the community (Fig. 2A). Based on this “trunk-branch” concept, we developed a strategy to construct similarity networks and define communities and the specific implementation process is as follows:

1. Communities with more than 20 nodes in the initial-threshold network are defined as core communities (trunks). The initial thresholds were set at e-value = 1e-68 for the SSN and Z-score = 45.9 for the StSN.
2. Thresholds were gradually relaxed to extend the branches of each core community. Newly appearing nodes were assigned to the community with which they shared the highest number of edges. The SSN threshold was relaxed in logarithmic steps (e.g., 1e-68, 1e-67, …), while the StSN threshold was relaxed in steps of 0.1 Z-score (e.g., 45.9, 45.8, …).
3. Communities were merged based on their inter-community connectivity. If the proportion of edges connecting two communities exceeded 50% for either community, they were merged. This process was repeated until all inter-community connectivities fell below the 50% threshold.
4. New communities with more than 20 unassigned nodes were identified and included.
5. Steps 2–4 were repeated until the network reached a stage of edge explosion and massive community merging.

Previously, final visualization thresholds for sequence similarity networks were often chosen based on known functional annotations. However, such information is lacking for most ORs. We found that under different thresholds, the modularity of community structures defined by the “trunk-branch” strategy showed a sharp inflection point when compared with theoretical modularity derived from the Louvain algorithm. Here, we propose a quantifiable approach for determining the final community display threshold by identifying the inflection point. Validation against known data confirmed that the resulting community partitions are highly reliable.

### Sequence similarity network

First, we performed pairwise sequence alignment of all ORs using MMseqs2(Steinegger and Soding 2017) to obtain sequence similarities and e-values. To improve network readability, we applied pre-clustering with CD-HIT(Li and Godzik 2006; Fu et al. 2012) v4.8.1 (‘-c 0.6 -n 4’) to reduce redundant edges between highly similar nodes, resulting in a total of 4,947 nodes. The edge weight between nodes was defined as the average e-value of all OR pairs they represent. We then constructed the sequence similarity network based on the “trunk-branch” strategy (see "Trunk-Branch Strategy"). An e-value threshold of 1e-25 was ultimately selected as the cutoff for defining the final sequence similarity network and OR subfamilies. At this threshold, 37 OR communities were identified, comprising 8,457 ORs. The network was visualized using the organic layout algorithm in Cytoscape(Shannon et al. 2003).

### Structure similarity network

We performed structural modeling of all ORs using AlphaFold2(Jumper et al. 2021). Pairwise structural comparisons were then conducted using Dali(Holm 2022) to obtain Z-scores. Following an approach similar to that used in SSN, we applied a greedy clustering algorithm to pre-cluster structurally similar ORs at a Z-score threshold of 50, resulting in 3,854 nodes. The edge weight between nodes was defined as the average Z-score of all OR pairs they represent. We then constructed the structure similarity network using the “trunk-branch” strategy. A Z-score threshold of 42.9 was selected as the final cutoff for defining the StSN and OR structural subfamilies. At this threshold, 27 OR structural communities were identified, comprising 7,750 ORs. The network was visualized using the organic layout algorithm in Cytoscape(Shannon et al. 2003).

### Phylogenetic tree and selection pressure analysis

To infer the amino acid phylogeny of ORs within StrC17, we performed multiple sequence alignment using MAFFT(Katoh and Standley 2013) v7.505 with the parameters ‘--globalpair -- maxiterate 1000’. The alignments were trimmed using trimAl(Capella-Gutierrez et al. 2009) v1.4.rev22 with the parameter ‘-automated1’. A maximum likelihood tree was then constructed using IQ-TREE(Minh et al. 2013; Nguyen et al. 2015) v 2.2.0 with the parameters ‘-m MFP -B 1000 --bnn’. To estimate dN/dS ratios, we used the codeml program from the PAML package (RRID:SCR_014932)(Yang 1997; Yang 2007). The codeml analysis employed the branch model with the control file parameters set to ‘model = 2’, ‘NSsites = 0’. We used a custom script to extract log-likelihood values (lnL) from the output files and perform likelihood ratio tests (LRTs).

### Evaluation of OR binding pocket properties

We used fpocket to identify OR binding pockets and assess their properties, with parameters set to ‘-m 3.1 -i 10’. To locate the centroid region of the binding pocket, we aligned each OR structure to MhOR5 using USalign(Zhang et al. 2022) v 20220924. Among all candidate pockets identified by fpocket(Le Guilloux et al. 2009) 4.0, the one closest to the centroid was designated as the binding pocket. Due to the stochastic nature of OR modeling, some ORs exhibited open binding pockets, making it difficult to define cavity-related parameters. To ensure consistency in the quantitative analysis, we only included ORs with closed binding pockets. Specifically, we retained only binding pockets with Number of Alpha Spheres < 140 and Volume < 800, excluding any that were open to the exterior.

### Molecular dynamics simulation of Orco β-Sheet structural function

We used GROMACS(Abraham et al. 2015) to simulate the interaction between the EL2 regions of OR and Orco with VOCs. The proteins used were the cryo-EM structures of ApOrco and ApOR5(Wang et al. 2024) (PDB: 8z9z), and the selected VOC was geranyl acetate (CID: 1549026). The membrane system was rapidly constructed using the QwikMD plugin in VMD(Humphrey et al. 1996). After default minimization and annealing steps, an optimized membrane-protein system was obtained. The Caver(Jurcik et al. 2018) tool was used to identify the pocket tunnels in the cryo-EM structures of ApisOR5 and ApisOrco. Surface residues were identified using FindSurfaceResidues(https://pymolwiki.org/index.php/FindSurfaceResidues), helping define the residues surrounding the binding pocket. Geranyl acetate was positioned above the pocket, and 5 ns MD simulations were performed to evaluate interactions.

A binding state was considered potential if the following criteria were met:

- The minimum distance between geranyl acetate atoms and pocket residue atoms was <12 Å (cutoff).
- The spatial distribution of the ligand satisfied the following exclusion conditions:
- Not located on the lateral side of the protein
- Not embedded in the phospholipid bilayer
- Not sterically blocked by the EL2 of ApisOrco

Each simulation was repeated 60 times for both ApOrco and ApOR5.

It should be noted that the molecular dynamics simulations were designed to examine the early trajectories of VOCs near the extracellular entrance of the binding pocket, rather than to assess the conformational stability of the receptor complex. A duration of 5 ns is sufficient for small molecules to respond to local steric constraints.

### Heterologous expression and calcium imaging

pDmelOR-mRho.V5.mER.hOr47a (RRID:Addgene_126478), a high-copy number bidirectional expression vector by inserting *D. melanogaster* Or47a (DmOr47a) and Orco in the pCMV-BI vector (RRID:Addgene_126475) backbone(Miazzi et al. 2019), was a gift from Bill Hansson (Addgene plasmid # 126478), then V5 tag was replaced with Flag tag, named pDmelOR-OR47a. The EL2 of OR47a (GHAEPELPFPCLFPWNIHI) in pDmelOR-OR47a was replaced by the EL2 of Orco (GDSVKMVVDHETNSSIPVEIPRLPIKSFYPWNASHG) to construct the new plasmid of pDmelOR-el2.OR47a. The sequences were confirmed by Sanger sequencing (Sangon Biotech, China).

HEK293T cells (RRID: CVCL_0063) were purchased from Wuhan Procell Biotechnology and maintained in DMEM medium (BasalMedia, China) supplemented with 10% fetal bovine serum (ExPhos, China) and 1% penicillin−streptomycin mixture (Sangon Biotech, China) at 37 °C with 5% CO_2_ until reaching 70–80% confluency. Cells were transfected with plasmids using LipoFiter transfection reagent (Hanbio, China). 6-8 h after transfection, transfected cells were seeded at a density of 6 × 10^5^ cells/well into a 35 mm glass-bottom culture dish (NEST, China). The HEK293T (RRID: CVCL_0063) cell line used in this study is derived from a male human fetus. HEK293T (RRID: CVCL_0063) cells were authenticated by STR profiling (Wuhan Procell) and tested negative for mycoplasma contamination every 3 months.

Cells were harvested for calcium imaging approximately 36 hours post-transfection. The culture medium was removed, and cells were rinsed with HBSS solution (BasalMedia, China). Subsequently, cells were incubated with Fluo-8 AM (5 μM, AAT Bioquest, USA) diluted in HBSS + 0.04% pluronic acid (F127, Sangon Biotech, China) in a CO_2_ cell culture incubator for 30 minutes at 37°C. Following three washes with HBSS solution, the Ca^2+^ dependent fluorescence was recorded with a fluorescence microscope (DeltaVision Ultra, GE) with the emission wavelength set at 525 nm and the excitation wavelength at 490 nm. Experiments lasted 5 min with a sampling interval of 5 s. The stimuli—pentyl acetate (PA, 100 μM; Aladdin, China), an OR47a agonist, and VUAA1 (100 μM; Sigma-Aldrich, USA), an Orco agonist—were gently added to the culture dish at 50 s and 180 s, respectively, after the start of the imaging protocol. The recorded signals were analyzed by Fiji (RRID:SCR_002285) ImageJ v.1.54 (RRID:SCR_003070) (Schindelin et al. 2012). Levels of Ca^2+^ were expressed as ΔF/F0 ratios. The average fluorescence intensity of the cells during the first 30 seconds was recorded as the baseline (F0). Each assay was conducted in triplicate to ensure reproducibility. For calcium imaging experiments, data acquisition and initial ΔF/F0 ratio calculations were performed by researchers who were blinded to the treatment groups during the analysis phase. Sample sizes for calcium imaging (n = 30) were determined based on common standards in the field and prior pilot experiments to ensure sufficient statistical power for detecting changes in fluorescence intensity

### Virtual docking data of ORs and VOCs

We constructed an insect-relevant odorant molecule library from four sources: pheromones(Li et al. 2023), food odors(Chi et al. 2025), fragrances(thegoodscentscompany.com), and theoretical odors(Mayhew et al. 2022). This approach ensured ecological relevance and chemical diversity. After deduplication, a total of 14,499 VOCs were retained for downstream analysis. We used Dragon (v.5.4)(Haddad et al. 2008) to calculate 32 representative molecular descriptors for each VOC, and the ChemmineR(Cao et al. 2008) R package to identify functional groups (Supplementary Data S8). VOCs were assigned functional group labels based on group priority rules. Dimensionality reduction of the 32 descriptors was performed using t-SNE, and the resulting two-dimensional odor space was visualized using Matplotlib(Hunter 2007). All VOC SMILES were converted to db2 format using the online resource tldr.docking.org. OR structures were aligned to the cryo-EM structure of MhOR5 (PDB:7LIC) to estimate the approximate position of the binding pocket. Virtual docking was then performed using DOCK 3.8 (RRID:SCR_000128)(Coleman et al. 2013), with batch processing handled via custom scripts ‘prepare_dock_file.sh’ and ‘run_dock_paraFly.sh’. In total, we obtained 130 million docking records between ORs and VOCs (Supplementary Data S9).

### Hit rate curve prediction and maximum hit estimation

We first compiled published functional datasets containing explicit OR-VOC response relationships. These datasets included ORs from *Drosophila melanogaster*, *Anopheles gambiae*, *Helicoverpa armigera*, *Spodoptera littoralis* and *Locusta migratoria*. For each dataset, OR-VOC combinations were assigned response or non-response labels according to the original experimental results. Each OR-VOC pair was then docked using the same structural modeling, docking, and scoring workflow used in the global OR-VOC analysis. Pairs were grouped into docking-score intervals, and the hit rate for each interval was defined as the proportion of experimentally positive pairs among all pairs in that interval. Each score interval was sampled three times. To determine the docking score range at which VOCs are likely to exhibit binding activity in electrophysiological assays, we applied Bayesian statistical modeling using prior probability distributions to define the top, midpoint, random hit rate, and slope of the hit rate curve. We assumed that the relationship between docking score (energy, ei) and hit rate follows a dose-response-like curve (fig. S10 A)(Lyu et al. 2019).

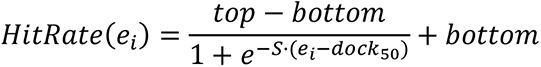

This function is defined by four parameters: (1) top: the maximum hit rate; (2) dock50: the docking score at which the hit rate is half of the top value (in kcal mol⁻¹); (3) S: the slope of the curve, calculated as S = slope × 4 / top, where slope is the percentage change in hit rate at dock50 per kcal mol⁻¹; (4) bottom: the minimum hit rate, fixed at 0. To define prior distributions, we analyzed OR hit rates for *Drosophila melanogaster*(Hallem and Carlson 2006), *Anopheles gambiae*(Carey et al. 2010), and *Spodoptera littoralis*(de Fouchier et al. 2017) across eight docking energy intervals and used this empirical data to derive posterior distributions of the model parameters. We specified independent Bayesian priors for each parameter: P(top) ∼ beta (α=20, β= 80), P(dock50) ∼ normal (μ = -10, σ = 15), P(S) ∼ normal (μ = -0.6, σ = 0.1) (fig. S10, B to D). Sampling from the prior distributions was performed using Hamiltonian Monte Carlo implemented in Stan, with parameters: ‘num_samples = 100000’, ‘num_chains = 4’, ‘num_warmup = 50000’, ‘max_depth = 12’.

### Docking-profile similarity network construction and community labeling

We first calculated the Pearson correlation coefficients (PCCs) for docking energies between all OR pairs across the VOC library. ORs were then ranked by the number of VOCs meeting the docking-score threshold, and a greedy clustering algorithm was applied to pre-cluster ORs with high docking-profile similarity (PCCs ≥ 0.6), resulting in 6,299 nodes. The edge weight between nodes was defined as the average PCC of all OR pairs. The docking-profile similarity network was then constructed using the “trunk-branch” strategy. Due to the complexity of OR functional divergence, our strategy yielded a modularity value that deviated by 0.1 from the theoretical Louvain modularity. To ensure accurate community assignment, we adopted the Louvain algorithm for final community detection. A PCC threshold of 0.48 was selected as the cutoff for defining the final docking-profile similarity network and OR docking-profile communities. The network was visualized using the organic layout algorithm in Cytoscape(Shannon et al. 2003).

Then, we assigned a VOC docking breadth label to each docking-profile community. First, we counted the number of VOCs with each functional group that were with docking scores below −8 kcal mol^-1^ in each docking-profile community. These counts were then compared against the total number of ORs recognizing those VOCs across all ORs, using Bonferroni-corrected post hoc tests. If a community showed a significantly higher or lower recognition count, it was considered to have a higher or lower docking-derived VOC count for that functional group. The results were normalized by the average VOC count to derive a docking-profile breadth score for each dPC. *Drosophila melanogaster* ORs were mapped to dPCs to evaluate the consistency between dPC labels and experimental response data. For each VOC functional group, ORs were assigned positive or negative labels according to their dPC labels, and these labels were compared with response counts from the Drosophila functional matrix to generate ROC curves and AUC values.

## Acknowledgments

We thank Yuan Huang from Shaanxi Normal University and Chenzhu Wang from the Institute of Zoology, Chinese Academy of Sciences, for their contributions to the early planning of the project. We thank Wenyu Zhang from Northwestern Polytechnical University and Yang Liu from the Chinese Academy of Agricultural Sciences for their suggestions on improving the conceptual framework during the mid-stage of the project. We also thank Hongbo Jiang and Fu Cao from Southwest University for their guidance on the experiments. We sincerely thank the reviewers for their constructive and thoughtful comments, which helped us improve the clarity, rigor, and presentation of this manuscript.

## Author contributions

Conceptualization: H.L., C.X.; Methodology: T.Z., X.Y., Y.F., Y.L.; Investigation: T.Z., X.Y., W.X., H.L., X.Y., Y.Z., S.D., C.G., Y.G.; Visualization: T.Z., X.Y., Y.Y.; Supervision: H.L., C.X., G.L.; Writing—original draft: T.Z., H.L., C.X., G.L.

## Data and resource availability

The data supporting the findings of this study (Supplementary Data S1–S9) are available for download from the figshare repository at https://doi.org/10.6084/m9.figshare.30541190.v3. All custom scripts and R code used for bioinformatics filtering and similarity network construction are available at GitHub (https://github.com/zhangtm0309/Code) and archived on Zenodo (DOI: 10.5281/zenodo.18459276).

