## supplementary information for "Class-wide sequence, structural, and docking-profile diversity of insect odorant receptors and structural specialization of Orco"

**This PDF file includes:**

Supplementary Text S1 and S2
Figs. S1 to S10

**Other Supplementary Materials for this manuscript include the following:**

Movie S1
Table S1 to S3
Datasets S1 to S9

### Supplementary Text S1

#### Benchmarking of OR annotation completeness and structural filtering

To evaluate the completeness and reliability of the odorant receptor (OR) annotation workflow, we benchmarked the automated annotation and structural-filtering pipeline against curated OR repertoires from three well-studied insect species: *Drosophila melanogaster*, *Anopheles gambiae*, and *Bombyx mori*. These species were selected because their OR repertoires have been extensively annotated and are among the best curated datasets available for comparison. The benchmark was designed to assess three aspects of the workflow: (i) recovery of previously reported ORs by the automated annotation pipeline, (ii) the effect of structural filtering on OR model quality, and (iii) whether candidate ORs in tandem-rich regions could reflect exon-merging artifacts.

For each benchmark species, we recorded the number of ORs recovered before structural filtering, the number retained after structural filtering, and the correspondence between retained candidates and literature-annotated ORs. Candidate ORs were matched to published ORs based on sequence similarity (Table S2). When discrepancies occurred, the corresponding sequences and predicted structures were manually inspected. For *Bombyx mori* candidates that did not correspond one-to-one to literature-annotated ORs, we further examined genomic locations to evaluate whether they may have resulted from exon merging among adjacent tandem ORs (Table S3). Candidates located on different chromosomes or outside the same tandem region as their closest literature ORs were not considered likely exon-merging artifacts.

For *D. melanogaster*, the accepted repertoire contains 60 OR genes and 65 protein products including splice variants. At the gene level, the automated workflow recovered most benchmark ORs before structural filtering, but Or47b and Or98b were not annotated. After structural filtering, fragmented or structurally incomplete ORs were removed, leaving 56 ORs that otherwise correspond one-to-one with the benchmark ORs. Two ORs that had been annotated before filtering, DmOr59a and DmOr85e, were removed by the structural filter because DmOr59a was fragmented and DmOr85e lacked TM7 in the predicted structural model.

The published *A. gambiae* OR repertoire contains 79 OR genes. The automated annotation workflow initially recovered 77 ORs, including 72 previously reported ORs. After removing fragmented ORs and five literature-annotated full-length ORs, 67 structurally intact ORs were retained. The retained ORs otherwise corresponded to literature annotations. Because five published full-length ORs were excluded by the structural filter, we further examined their predicted structures. AgOr52, AgOr47, and AgOr64 lacked one complete transmembrane region. AgOr58 and AgOr6 contained all expected transmembrane regions, but showed local deletions in part of the transmembrane regions. These results indicate that structural filtering can remove clearly incomplete ORs, but may also exclude a small number of genuine ORs with atypical or locally incomplete transmembrane-region features.

The published *B. mori* OR repertoire contains 66 OR genes, including two pseudogenes. The automated workflow initially annotated 103 OR candidates, including 60 literature-annotated ORs. After structural filtering, fragmented ORs and seven literature-annotated ORs were removed, and 64 structurally intact ORs were retained. Among these, 53 corresponded to literature-annotated ORs. Some retained candidates showed high sequence similarity to literature ORs, whereas others showed lower similarity and did not clearly correspond to previously reported ORs. To assess whether such candidates could result from exon-merging errors in tandem OR regions, we examined their genomic positions. Most candidates did not occur on the same chromosome or in the same tandem region as their closest literature-annotated ORs. Therefore, these candidates are unlikely to be simple exon-merging artifacts involving adjacent OR genes, and may represent previously unannotated OR candidates. Nevertheless, transcriptomic evidence or manual gene-model curation will be needed to confirm these candidates.

Because previously known OR sequences from some benchmark species were included in the query database, we next tested whether the presence of conspecific queries systematically influenced the final inferred OR repertoire size. We divided the benchmark species into two groups: those with conspecific OR sequences represented in the query database and those without conspecific queries. We detected no significant difference in OR annotation performance between the two groups ($P = 0.8$, Fig. S1B). Because *Anagrus nilaparvatae* showed an unusually large difference in OR number between the two studies, we further performed a sensitivity analysis excluding this species, and the conclusion remained unchanged ($P = 0.4$, Fig. S1C). Therefore, among species for which comparable genome-based reference repertoires are currently available, we found no evidence that the presence or absence of conspecific ORs in the query database systematically affects the final inferred OR repertoire size.

To extend the benchmarking beyond these three detailed case studies, we compared the final numbers of structurally intact ORs with published repertoire counts from additional insect species (Table S1). For most species with comparable genome-based reference repertoires, the number of ORs retained in this study exceeded 70% of the published count and showed generally good agreement in repertoire size. *Diabrotica virgifera virgifera* was an exception: a previous manual annotation reported 124 complete ORs, whereas our pipeline retained 78 structurally intact ORs, approximately 63% of the published count. This discrepancy likely reflects the challenges posed by the large, repeat-rich genome of *D. virgifera virgifera*, in which repetitive sequences and complex OR loci can complicate automated gene prediction and locus reconstruction. The previous species-specific annotation incorporated transcriptomic evidence and locus-by-locus manual curation, which can help resolve gene models in complex genomic regions. Therefore, transcriptome-assisted manual annotation may be required to obtain more comprehensive OR repertoires for species with large, repeat-rich, or otherwise complex genomes.

Overall, the benchmarking analysis indicates that the automated annotation workflow recovers most known OR repertoires in well-annotated species and that structural filtering improves dataset reliability for downstream structural comparisons and docking-profile analyses. The final dataset was deliberately designed as a conservative, high-confidence set of structurally intact ORs rather than the broadest possible species-level gene catalog. Accordingly, lower counts relative to some published repertoires are an expected consequence of uniformly excluding fragments, pseudogenes, and candidates that do not meet the structural inclusion criteria. To evaluate whether this conservative filtering influenced the ecological analyses, we repeated the phylogenetic comparative analyses using the larger of our structurally intact OR count and the corresponding published count. The main associations between OR repertoire size and ecological traits remained unchanged (fig. S1D–G), indicating that the filtering strategy did not affect our conclusions.

### Supplementary Text S2

#### Hit-rate benchmark for docking-score threshold selection

To evaluate whether the docking-score threshold was driven by any single functional dataset, we performed hit-rate analyses using experimentally characterized OR-VOC pairs from multiple insect groups.

When *Drosophila melanogaster, Anopheles gambiae*, and Lepidoptera datasets were analyzed separately, they showed similar docking score-hit rate relationships. In all three datasets, experimentally responsive OR-VOC pairs were enriched as docking scores became more negative. The hit-rate plateau for these datasets was approximately 23%, and the corresponding docking-score threshold was around -8 kcal/mol. This result is consistent with the threshold used in the main analysis and indicates that the -8 kcal/mol cutoff was not driven by a single species or dataset.

We also included *Locusta migratoria* OR functional data as an independent sensitivity analysis. The locust dataset differed from the other datasets in two important ways. First, the experimental positive rate was low, approximately 5.2%. Second, previously characterized locust ORs are mostly narrowly tuned receptors.

Consistent with these properties, the locust dataset showed lower overall hit rates than the Drosophila, Anopheles, and Lepidoptera datasets. In most docking-score intervals, the locust hit rate remained below 10%. When the locust dataset was combined with the other three datasets, the overall hit-rate plateau decreased to approximately 18%, and the corresponding docking-score threshold shifted to approximately -11 kcal/mol.

The locust result indicates that OR tuning breadth and the experimental positive rate of a dataset can influence the relationship between docking score and hit rate. For datasets dominated by narrowly tuned ORs and low experimental positive rates, a stricter docking-score threshold may be required to achieve stronger enrichment of experimentally responsive OR-VOC pairs.

However, the goal of the present study was to establish a unified empirical threshold for large-scale comparison across thousands of ORs and a broad VOC library, rather than to optimize separate thresholds for individual lineages. Therefore, the -8 kcal/mol threshold derived from the Drosophila, Anopheles, and Lepidoptera datasets was retained for the main analysis, whereas the locust result was treated as a sensitivity analysis and used to define the limitation of applying a single cutoff across all insect groups.

The hit-rate benchmark supports the use of docking scores for repertoire-level enrichment and comparative analysis. Nevertheless, the plateau hit rate remains limited and is not sufficient to assign definitive ligands or experimentally validated response spectra to individual ORs.

Therefore, docking results in this study should be interpreted as docking-derived binding potential. They provide a high-throughput, internally comparable framework for identifying repertoire-level trends and candidate OR-VOC relationships for future testing, but specific receptor-ligand pairs require validation by heterologous expression, electrophysiology, calcium imaging, genetic perturbation, or behavioral assays.

### Figures

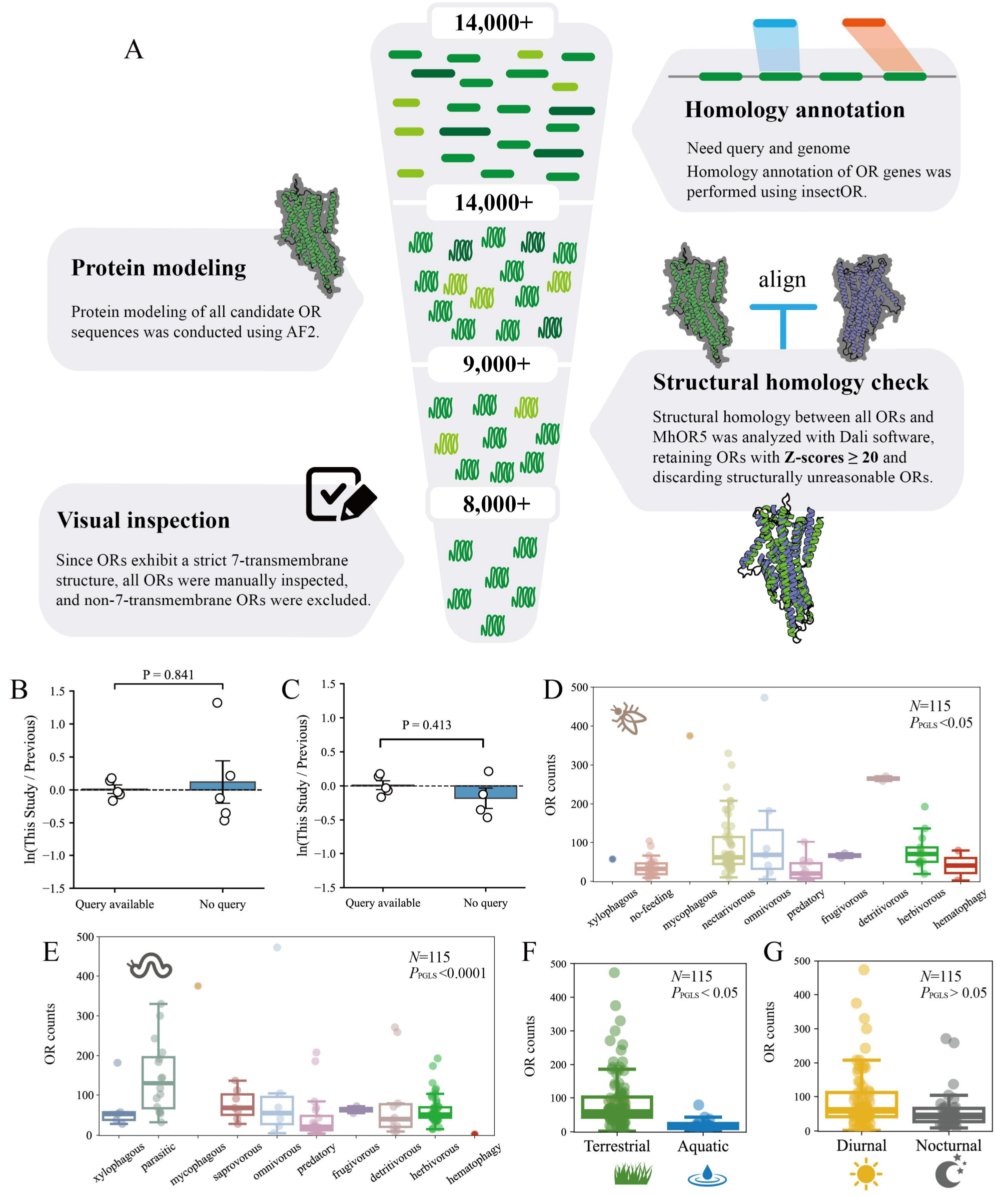


**Fig. S1. Structural screening strategy and pGLS sensitivity analysis of OR quantity** (A). The stringent screening strategy for complete OR genes. All homologously annotated OR genes underwent structure modeling. Structural similarity was used as an indirect measure to assess sequence reliability (Dali Z-score > 20). Finally, manual inspection was performed to refine the intact OR genes. For detailed procedures, refer to the Methods. (B) Comparison of OR counts from this study and previous annotations between species with and without conspecific sequences in the query set. Significance was assessed using the Mann–Whitney U test. (C) Comparison after excluding the OR annotation results for the outlier *Anagrus nilaparvatae*. (D-G). pGLS tests for adult diet, larval diet, habitat, and circadian rhythm after OR counts were updated to the larger of structurally intact OR counts and published OR counts.

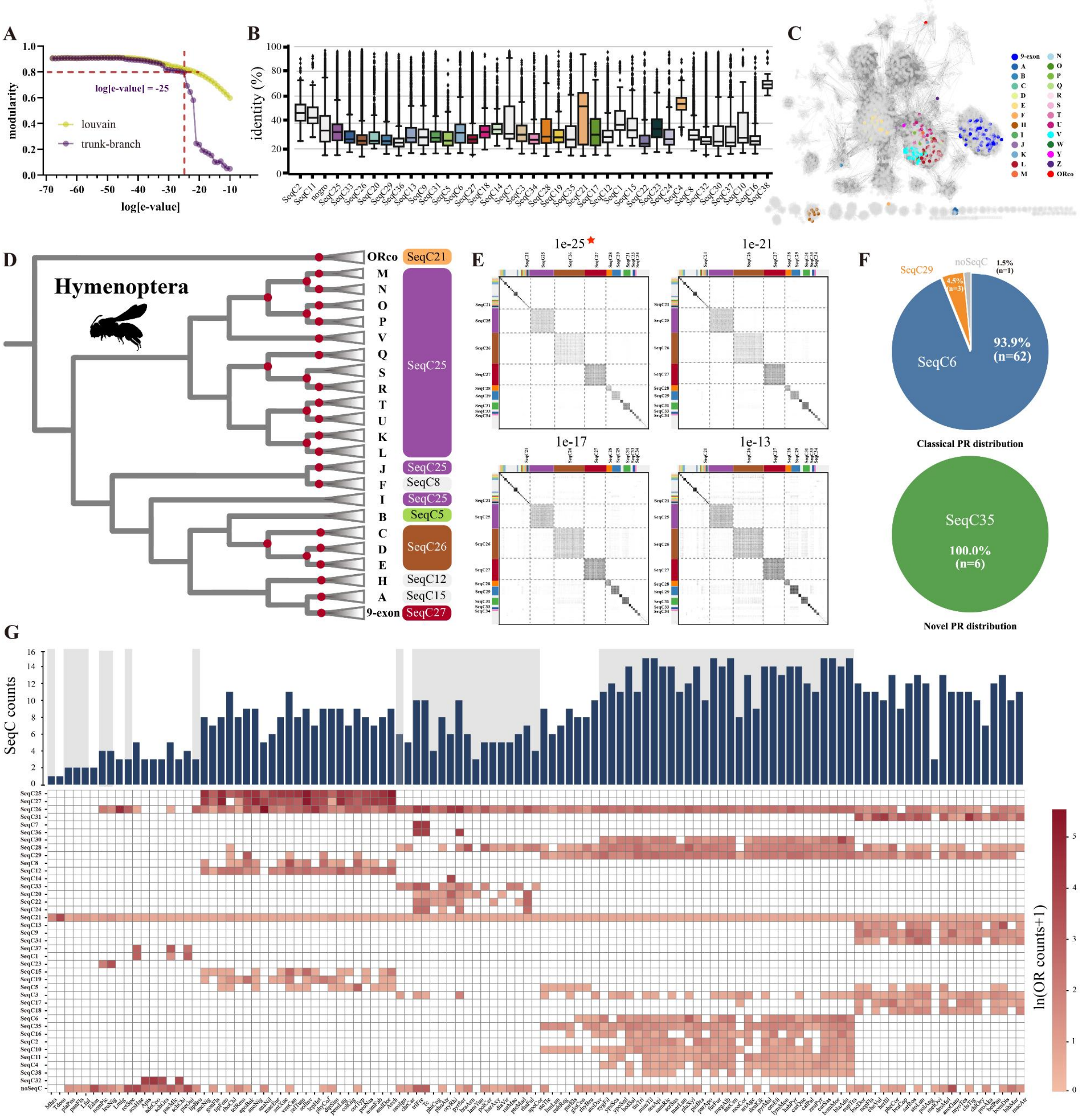


**Fig. S2. SSN intra-subfamily OR sequence similarity and evidence supporting the accuracy of SeqC.** (A). Modularity changes between SeqCs at different SSN alignment e-value thresholds. Purple represents modularity obtained using the "trunk–branch" algorithm, while yellow denotes the theoretical optimal modularity based on the Louvain algorithm. (B). Pairwise sequence similarity within different SeqCs. The SeqC colors are consistent with Figure 2C, while gray indicates SeqCs with no interactions with others. Boxplots represent the first quartile minus 1.5 times the interquartile range, the interquartile range, the first quartile, the mean, the third quartile, and the third quartile +1.5 interquartile range. Dots indicate outliers. (C). Mapping of Acromyrmex echinatior (Aech) OR subfamilies onto the SSN. Different colors represent previously defined hymenopteran subfamilies. (D). Phylogenetic relationship between previously defined hymenopteran subfamilies and our SeqC classification. Red dots indicate branches with bootstrap values greater than 80. Branches with the same subfamily designation as previous studies are collapsed. The phylogenetic tree is based on the previous study24. Animal silhouettes were obtained from PhyloPic.org. (E). Greyscale interaction matrix showing interactions between SeqCs under different e-value thresholds. At an e-value of 1e-25, SeqC boundaries are distinct. As the threshold increases to 1e-17, interactions between SeqCs become more frequent, with extensive interactions observed at 1e-13. The SeqC divisions remain clear within approximately 8 orders of magnitude. Different color bars represent different SeqCs, consistent with Figure 2C. (F). Proportions of Lepidopteran PRs distributed across different SeqCs. (G). Heatmap showing the number of ORs within different SeqCs across species. Due to significant variation in OR numbers among species, the OR counts are ln-transformed. The top bar plot represents the number of SeqC types in each species. Gray shading indicates boundaries between insect orders.

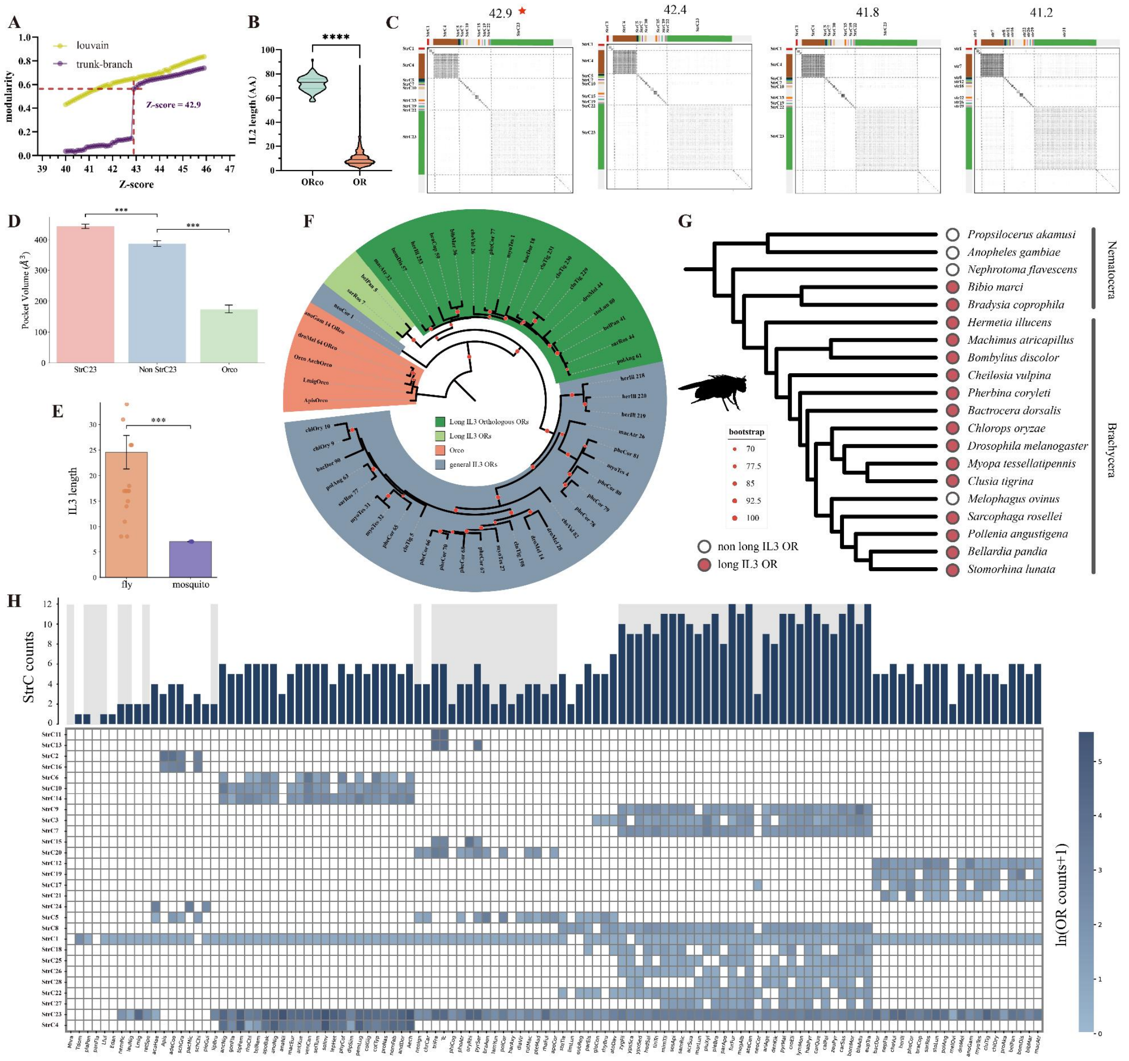


**Fig. S3. StrC classification and the phylogenetic tree of StrC17 ORs.** (A). Modularity changes between StrCs at different Dali Z-score thresholds. Purple represents modularity obtained using the "trunk–branch" algorithm, while yellow denotes the theoretical optimal modularity based on the Louvain algorithm. (B). Comparison of IL2 loop amino acid lengths between all ORs and Orco. A two-sided t-test was used to assess significance. (C). Greyscale interaction matrix showing interactions between StrCs under different Z-score thresholds. At a Z-score of 42.9, the boundaries between StrCs are distinct. At a threshold of 41.8, interactions between StrCs increase, with extensive interactions observed at 41.2. Different color bars represent distinct StrCs, consistent with Figure 3B. Gray indicates StrCs without interactions. (D) Binding-pocket volume of the shared structural cluster StrC23. Tukey's post hoc test was performed after one-way ANOVA. (E) Comparison of IL3 length between the conserved fly OR cluster that recognizes 1-octen-3-ol and mosquito ORs that recognize 1-octen-3-ol. Welch's t-test was used. ***p < 0.001. (F). Maximum likelihood phylogenetic tree of StrC17 ORs, with Orco as the outgroup. Green indicates long IL3 ORs, and red circles mark bootstrap values. The tree was visualized using iTOL. (G). Species tree of dipteran species included in the study. Red circles represent species with long IL3 ORs, while hollow circles indicate species without long IL3 ORs. Animal silhouettes were obtained from PhyloPic.org. (H). Heatmap showing the number of ORs within different StrCs across species. Due to significant variation in OR numbers among species, the OR counts are ln-transformed. The top bar plot indicates the number of StrC types in each species. Shading marks the boundaries between insect orders. ****$P < 0.0001$

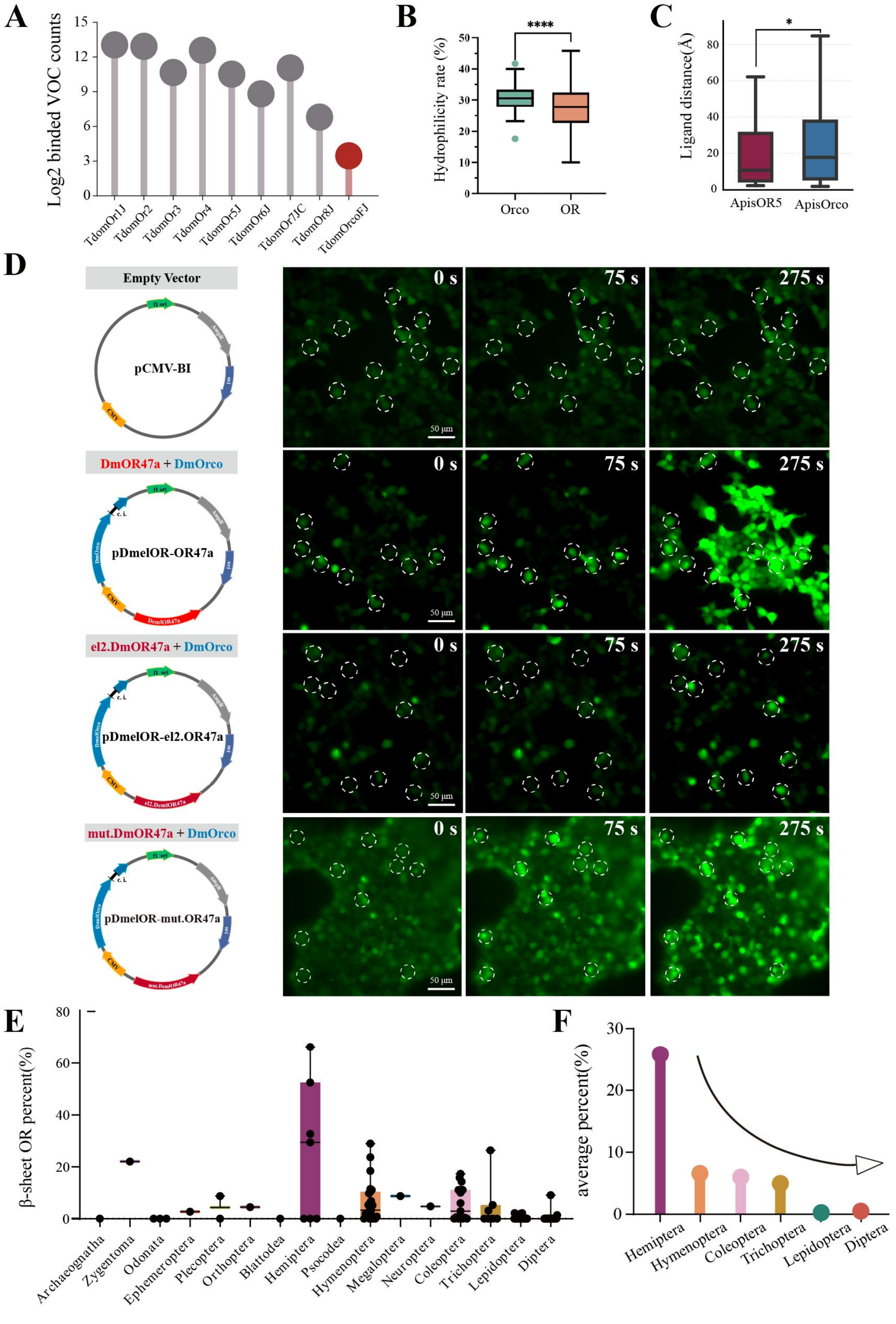


**Fig. S4. The driving factors underlying the origin of Orco.** (A). The number of VOCs recognized by Tdom1-8 and Orco. VOCs with a DOCK score < -8 kcal/mol were considered bound. The number of bound VOCs is shown on a log2 scale. (B). Proportion of hydrophilic amino acids in the EL2 region of Orco and OR. Statistical significance was assessed using a two-sided t-test. Dots represent outliers. (C). The distance between geranyl acetate and its initial position above ApisOR5 and ApisOrco was measured after MD simulation. The results indicate that Orco exhibits a dispersive effect on small molecules. (D). Examples of calcium imaging experiments with HEK293 cells co-transfected with DmOr47a + Orco constructs, the el2.DmOr47a mutant construct (in which the EL2 region of DmOr47a was replaced with that of Orco), and the mut.DmOr47a mutant construct (in which a short peptide is inserted into the EL2 region of DmOr47a at the position corresponding to the EL2 structure of Orco, rather than forming a β-sheet structure). Schematic diagrams of different plasmid constructs are shown on the far left. CMV: cytomegalovirus promoter. c.i.: chimeric intron. ori: origin of replication. Fluorescence changes are shown following stimulation with PA (middle column) and VUAA1 (right column). White dashed circles indicate the cells used for fluorescence quantification. The experiment was independently replicated three times; results from one representative trial are shown here. (E). Box plot showing the proportion of EL2 β-sheet ORs among total ORs for different insect orders. Dots represent the proportion of β-sheet ORs in each species. (F). Average proportion of β-sheet ORs among total ORs for six major insect orders. The proportion of β-sheet ORs decreases progressively with evolutionary time. * $P < 0.05$, **** $P < 0.0001$.

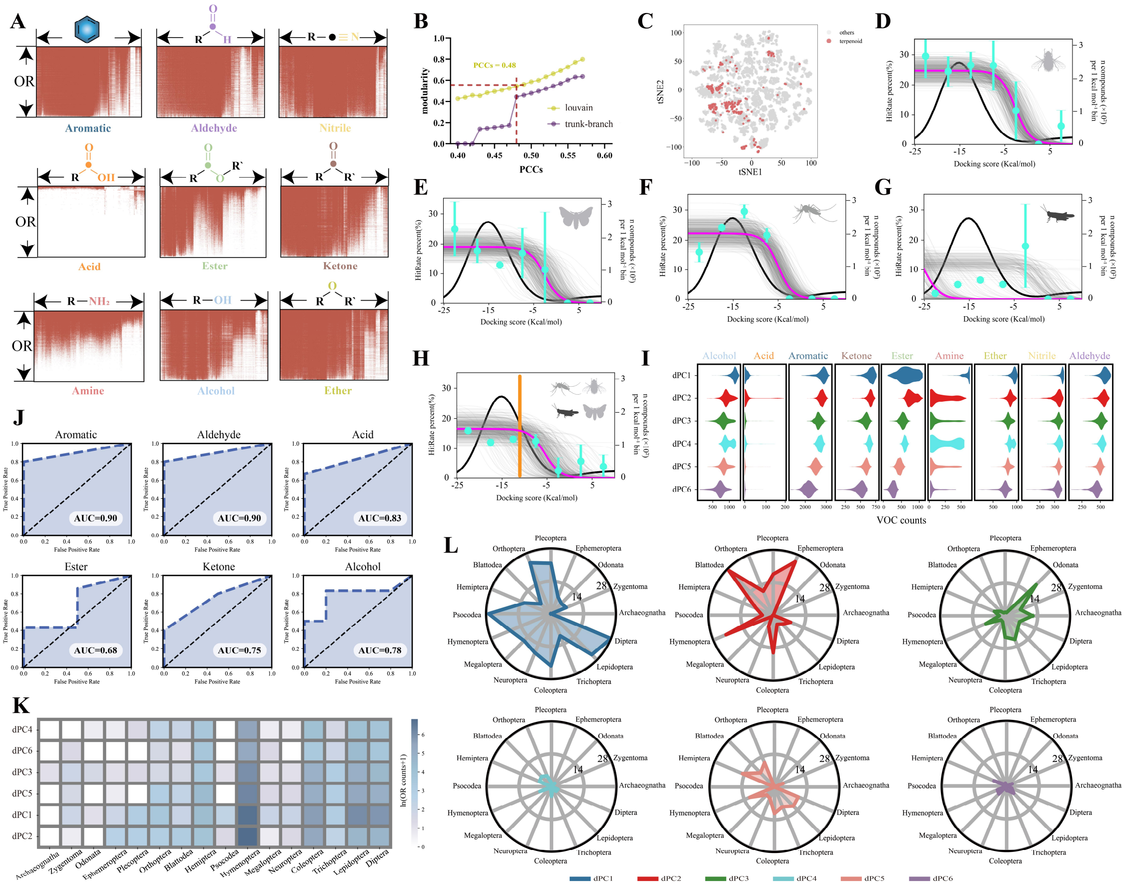


**Fig. S5. Insect OR-VOC binding profiles, dPC labeling accuracy, and distribution of docking-profile communities across insect orders.** (A). Binding profiles of insect ORs with VOCs from different functional groups. Each point represents the interaction between an OR and a small molecule. A docking score below −8 kcal $mol^{-1}$ was classified as predicted binding. (B). Modularity changes between docking-profile communities at different PCC thresholds. Purple represents the docking-profile community modularity obtained using the "trunk-branch" algorithm, while yellow denotes the theoretical optimal community modularity based on the Louvain algorithm. (C) Distribution of terpenoids in the two-dimensional odor space. (D-H) Hit rates for ORs from single insect taxa. Species include *Drosophila melanogaster*, *Anopheles gambiae*, *Helicoverpa armigera*, *Spodoptera littoralis*, and *Locusta migratoria*. Cyan dots represent the mean hit rate +/- s.e.m. for each docking-score interval. The gray curve represents posterior distributions from Bayesian inference, with n = 500. The black curve shows the distribution of hit rates and docking scores across all OR-VOC docking results. (I). Number of VOCs recognized by different dPCs. Colors represent different dPCs, consistent with Figure 5c. (J). Accuracy of dPC labeling using Drosophila ORs as an example. The AUC represents the relationship between the predicted small molecule recognition tendency labels and the experimentally validated number of recognized VOCs. (K). Number of ORs in different docking-profile communities across insect orders. Due to large variations in OR numbers among orders, the OR counts are ln-transformed. (L). Wilson scores for different dPCs across insect orders. Wilson scoring was applied to correct for potential errors caused by low OR numbers in certain orders, providing a more balanced evaluation of dPC representation.

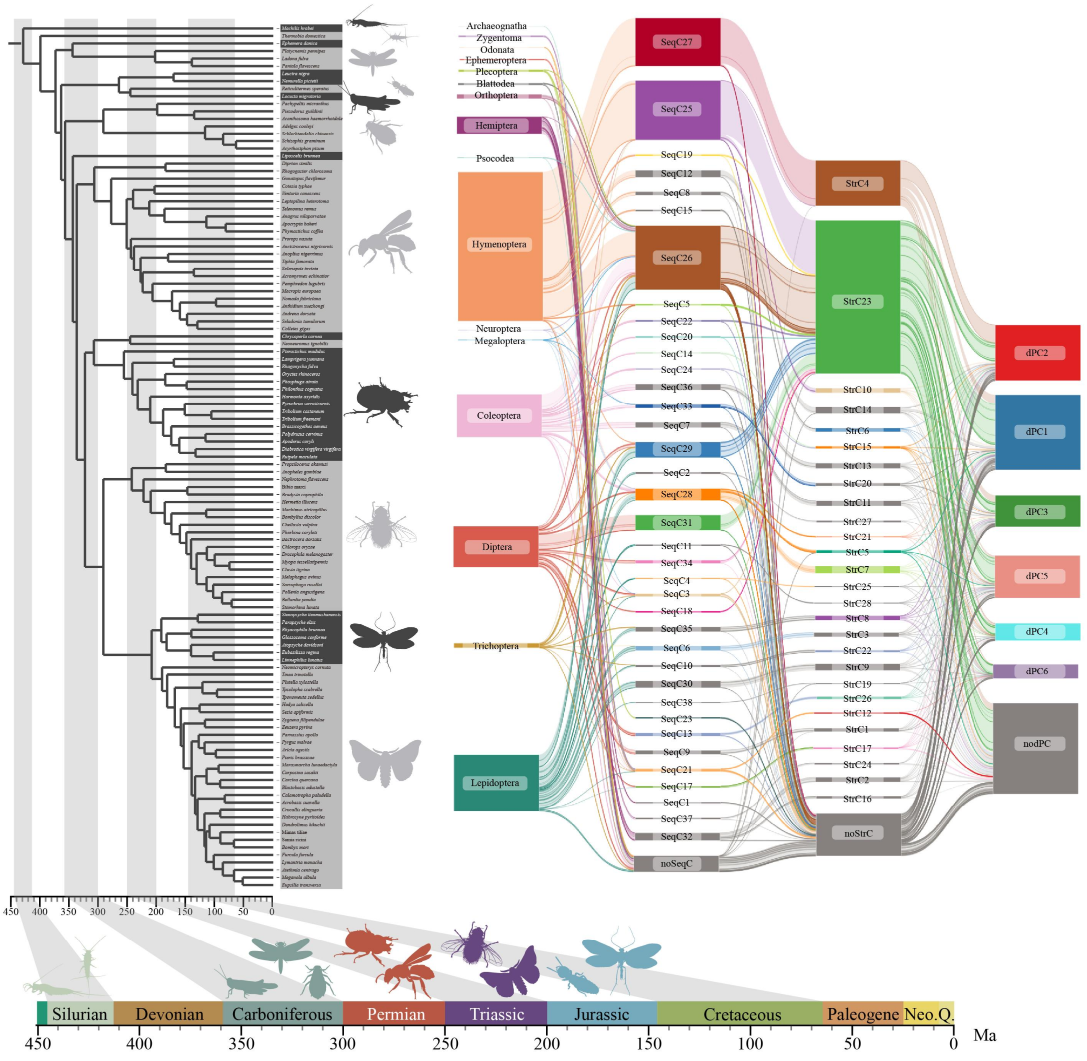


**Fig. S6. Relationships between OR sequence structure and docking-profile communities across insect orders (all ORs).** The phylogenetic tree follows the structure from Fig. 1. On the right, the mapping relationships between all insect OR SeqCs, StrCs, and dPCs are shown. Time nodes for different geological periods are based on previous studies1 and are separated by gray shading. The timeline beneath the tree marks the origin of various insect orders. Animal silhouettes were obtained from PhyloPic.org.

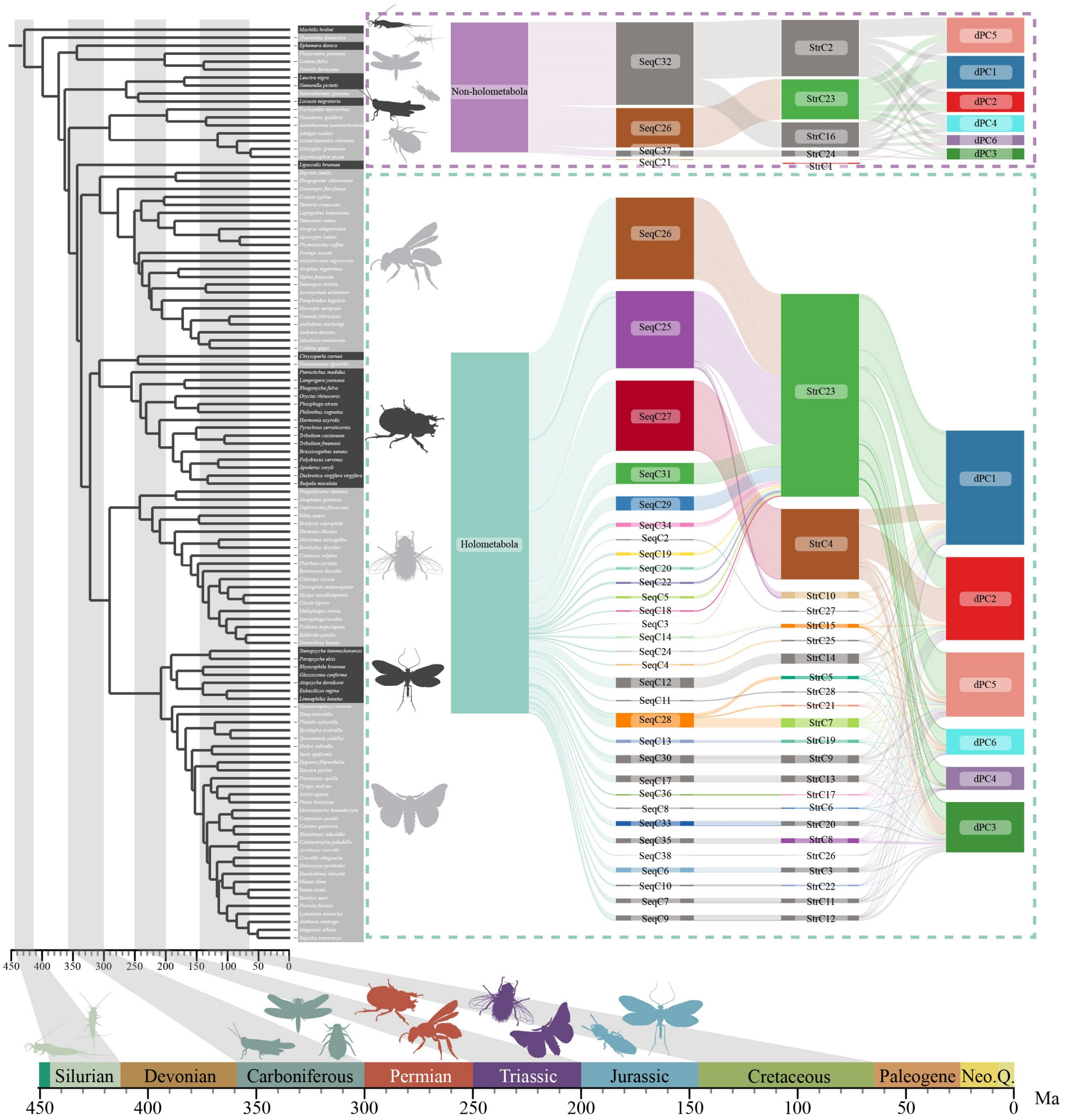


**Fig. S7. Sequence, structural, and docking-profile community associations of ORs in holometabolous and non-holometabolous insects.** The phylogenetic tree follows the structure from Figure 1. On the right, the mapping relationships between all insect OR SeqCs, StrCs, and dPCs are shown, considering only ORs classified into specific communities. Time nodes for different geological periods are based on previous studies1 and are separated by gray shading. The timeline beneath the tree marks the origin of various insect orders. Animal silhouettes were obtained from PhyloPic.org.

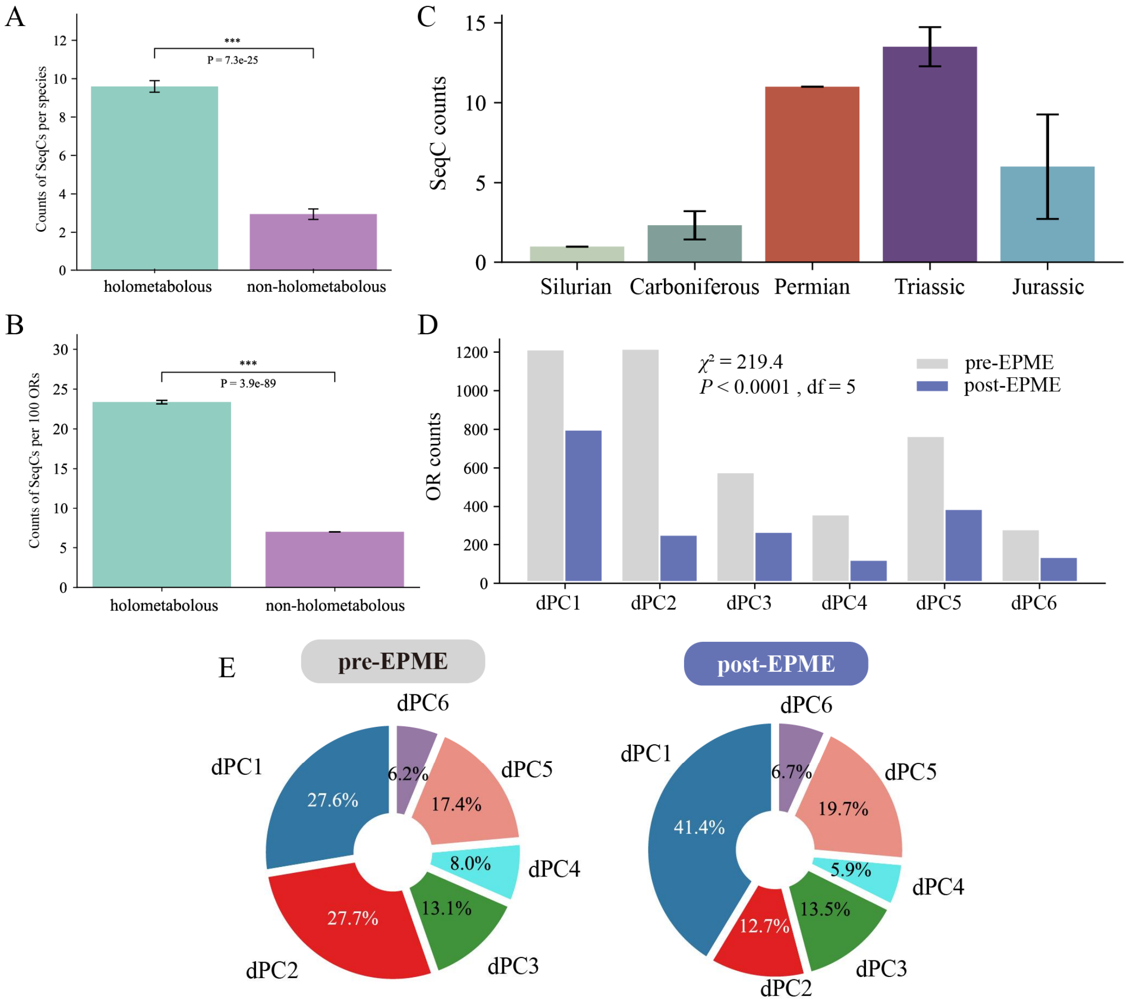


**Fig. S8. Changes in insect OR SeqC types and dPC proportions across different geological periods.** (A-B) Normalized analysis of SeqC diversity in holometabolous and non-holometabolous insects at the species level (A) and OR-count level (B). Welch's t-test was used. ***p < 0.001. (C). The number of SeqC types in insect ORs across different geological periods. (D). Changes in the number of ORs within different dPC categories before and after the EPME. (E). Proportional changes in different dPCs of insect ORs before and after the EPME. EPME: End-Permian Mass Extinction.

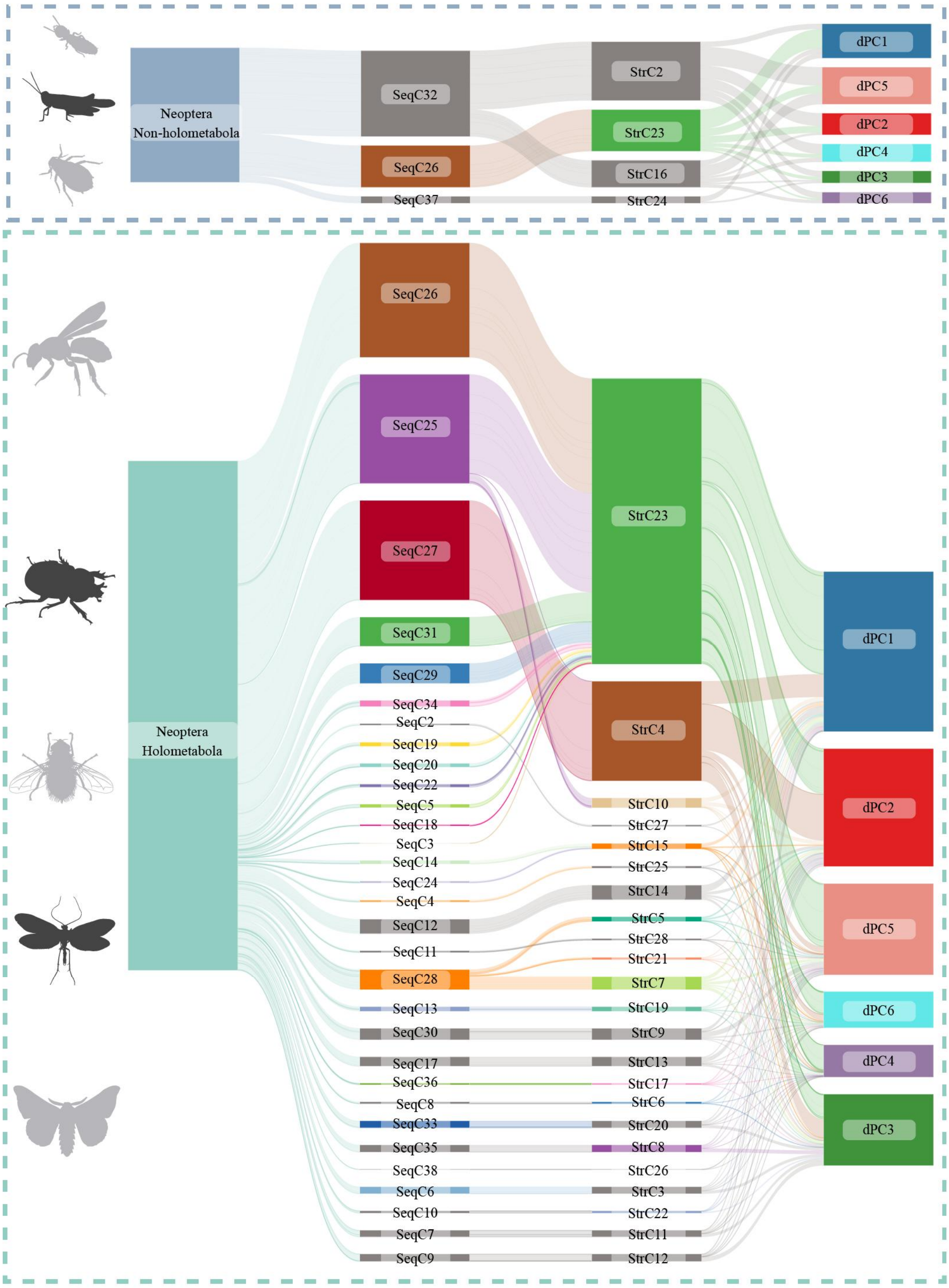


**Fig. S9. Comparison of OR SeqC diversity between Holometabola and non-holometabolous Neoptera.**

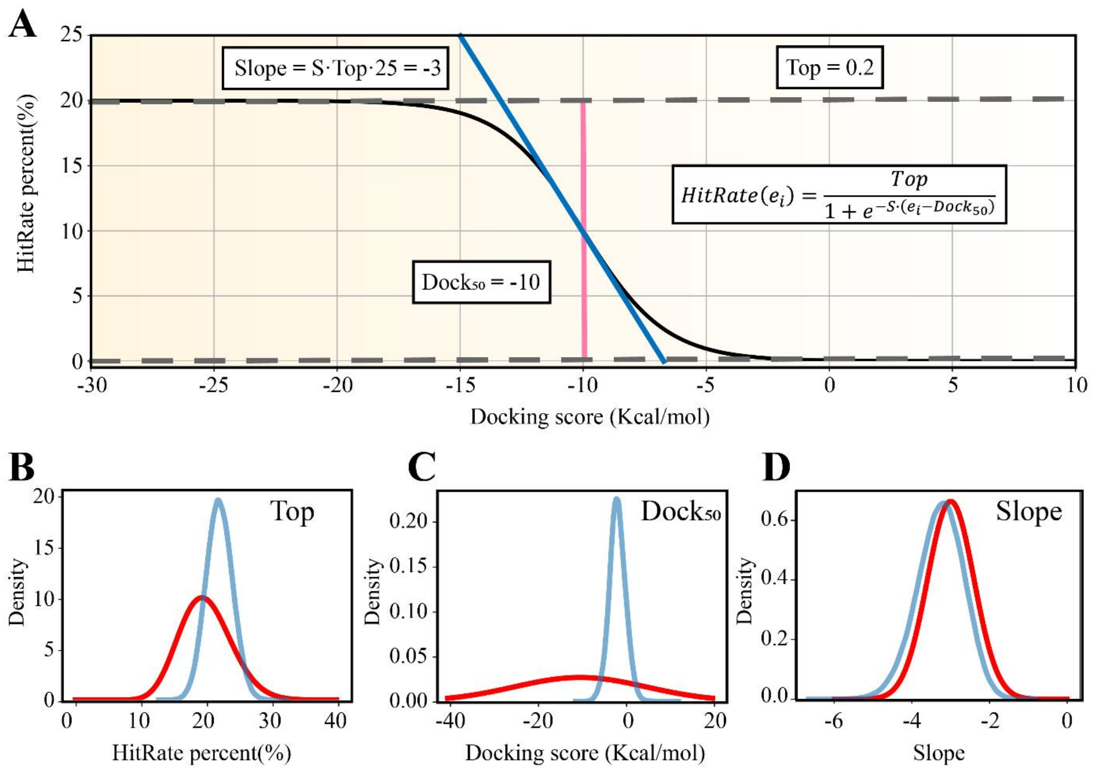


**Fig. S10. Bayesian prior modeling to optimize VOC selection by balancing information acquisition and ligand discovery.** (A). Sigmoid functional form for the hit-rate model. (B-D). Marginal Bayesian prior (blue) and posterior (red) distributions (n = 100,000) for each model parameter. B, Top. C, Dock50. D, Slope.

**Movie S1 (separate file).** Molecular dynamics trajectories of VOCs above the binding pockets of OR and Orco.

**Tables S1-S3 (separate file).** OR benchmark test

**Dataset S1 (separate file).** Genome information.

**Dataset S2 (separate file).** Amino acid sequences of insect ORs.

**Dataset S3 (separate file).** Living habits of 115 species.

**Dataset S4 (separate file).** Insect species tree.

**Dataset S5 (separate file).** Community annotation of insect ORs.

**Dataset S6 (separate file).** Orthologous groups identified within SeqC26, SeqC28, and StrC23

**Dataset S7 (separate file).** The likelihood ratio test results for StrC17 ORs.

**Dataset S8 (separate file).** The physicochemical properties and functional group labels of the 14,499 VOCs.

**Dataset S9 (separate file).** Matrix of all OR–VOC docking scores.
